# Single molecule imaging of transcription dynamics, RNA localization and fate in human T cells

**DOI:** 10.1101/2024.11.11.622939

**Authors:** M. Valeria Lattanzio, Nikolina Šoštarić, Nandhini Kanagasabesan, Branka Popović, Antonia Bradarić, Leyma Wardak, Aurélie Guislain, Philipp Savakis, Evelina Tutucci, Monika C. Wolkers

## Abstract

T cells are critical effector cells against infections and malignancies. To achieve this, they produce pro-inflammatory cytokines, including IFN-γ and TNF. Cytokine production is a tightly regulated process. The relative contribution of transcriptional and post-transcriptional regulation to mRNA expression is, however, unknown. We therefore optimized single-molecule FISH for primary human T cells (T-cell smFISH) to simultaneously quantify nascent RNA, mature mRNA levels and its localization with single-cell resolution. T-cell smFISH uncovered heterogeneous cytokine mRNA levels, with high cytokine producers displaying biallelic *IFNG*/*TNF* RNA transcription activity. Throughout activation, nuclear cytokine mRNAs accumulated, whereas cytoplasmic cytokine mRNA was degraded through translation-dependent decay. Lastly, T-cell smFISH uncovered cytokine-specific regulation by the RNA-binding protein HuR. Thus, T-cell smFISH provides novel insights in the intricate (post)-transcriptional processes in T cells.

## Introduction

CD8^+^ T cells play a crucial role in clearing infected and malignant cells. Upon target cell recognition, effector CD8^+^ T cells undergo a substantial remodelling of their transcriptome and proteome (*1–4*). This remodelling allows for the rapid release of effector molecules such as granzymes and pro-inflammatory cytokines (*5*, *6*). Key pro-inflammatory cytokines are IFN-γ and TNF, which license T cells to kill their target cells (*7–9*).

Cytokine production is regulated at multiple molecular levels, including transcriptional and epigenetic mechanisms (*10*, *11*). Another important layer of gene expression control is post- transcriptional regulation (PTR), which includes processes – such as RNA splicing, RNA transport, RNA stability, and translation control. The importance of PTR is evidenced by the fact that measured mRNA levels in T cells only mildly correlate with protein abundance (*4*), which is a highly conserved feature throughout evolution (*12*, *13*). For instance, memory T cells contain ready-to-deploy cytokine mRNA, which is blocked through translational control, yet becomes released from this block to serve as template for translation for rapid recall responses (*14*). Furthermore, cytokine overproduction of effector T cells induces immunopathology and autoimmunity, and loss of production results in dysfunctional anti-tumor responses (*8*, *15*). It is therefore of utmost importance to decipher how PTR processes control T cell function.

RNA binding proteins (RBPs) are key mediators of PTR that define the abundance of mRNA and protein in T cells (*6*, *16*, *17*). For instance, the RBP ELAVL1/HuR promotes early cytokine production of activated human CD8^+^ T cells (*6*). In other cell types, HuR was shown to mediate pre-mRNA splicing, control poly-adenylation, regulate the nuclear export of mRNA, and when located in the cytoplasm, promote mRNA stability and translation (*18–22*). Provided that RBP expression and their mode of action is cell-type specific (*23*, *24*), it remains to be uncovered how HuR promotes cytokine production in T cells.

The processing of an RNA molecule occurs at different subcellular locations (*25*). In the nucleus, the RNA is transcribed, spliced, and modified. When translocated into the cytoplasm, translation can take place. Previous reports that studied the regulation of mRNA expression in T cells measured mRNA transcription, mRNA kinetics, and their subcellular localization by bulk RNA- sequencing (*26*, *27*). However, these studies lack subcellular and single-cell resolution, which is crucial to understand the role of posttranscriptional gene regulation during T cell responses, given their inherently high heterogeneity (*28*, *29*). Single-molecule fluorescence in situ hybridization (smFISH) provides such in-depth insights (*30*, *31*). smFISH has been employed to visualize and quantify *-* with single-cell resolution - nascent RNAs at the transcription sites (*32*, *33*), quantify mature mRNA (*34–36*), or to identify its subcellular localization (*37*, *38*). While highly informative, these studies did not simultaneously measure *de novo* transcription the level and fate of mRNA in T cells. However, for deciphering gene regulation, integrating this information is paramount. Moreover, studies using smFISH in T cells (*32*, *39*) generally employ standard smFISH analysis pipelines that were developed for adherent cells (*40*, *41*). As these analysis pipelines are not geared towards the small cell size and round morphology of T cells, they fail to unequivocally call the subcellular localization of transcripts. To address these challenges and to comprehensively study cytokine RNA dynamics from transcription to translation, we optimized smFISH for T cells (T-cell smFISH). This optimized analysis pipeline enables the simultaneous quantification of nascent RNA, mature mRNA and the localization of nuclear and cytoplasmic mRNA, overcoming the limitations of the T cell morphology. Using this pipeline, we achieved an unprecedented depth of analyzing the fate of mRNA in T cells, here exemplified for cytokine mRNAs.

## Results

### 3D Quantification of single cytokine mRNA in primary T cells

To quantify cytokine mRNAs in human T cells, we first optimized the smFISH hybridization and analysis protocol (*see Materials and Method*s). To this end, we generated effector T cells (Teff) that can rapidly respond to recall responses, by activating human blood-derived CD8^+^ T cells for 72 h with α-CD3/α-CD28, followed by a 7-day of rest. Teff cells were subsequently restimulated with α-CD3/α-CD28, fixed, and simultaneously probed for *IFNG* and *TNF* mRNA by two-color smFISH (**Fig 1A**) (*31*, *42*). Using a wide-field fluorescence microscope, Z-stacks were acquired every 200 nm to encompass the entire cell thickness, and to precisely identify the *x*, *y*, *z* position of an mRNA (*43*). T cell outline was deduced from background-fluorescence in the smFISH channel and from differential interference contrast images (DIC; **Suppl. Fig. 1A**). DAPI was used as proxy for nuclear staining (**Suppl. Fig. 1A**). FISH-quant was used to enumerate total mRNAs (**Fig. 1B, 1C**). To identify the transcription sites (TsX), high-intensities signal (1.5x the average intensity of a single mRNA) colocalizing with DAPI staining was used (*see Materials and Methods)*. This approach allowed us to measure cells with one active TsX (mono-allelic) or two active TsX (bi-allelic) (**Supp. Fig. 1B**). The average intensity of all mRNAs was used as a reference to quantify the number of nascent mRNA per TsX. To localize individual cytokine mRNA molecules with subcellular resolution, we developed the T-cell smFISH pipeline, which is optimized to account for the small and compact structure of T cells by combining FISH-quant (*41*) for RNA quantification, filtering for cell selection, and Cell-Pose (*44*) for 3D mask reconstruction (**Suppl. Fig. 1C, Movie S1-S2;** https://github.com/nikolinasostaric/T-cell_smFISH). T-cell smFISH achieved a 98% accuracy for localization, as defined by manual analysis of 100 randomly chosen mRNAs. T-cell smFISH thus allowed us to elucidate the expression of cytokine transcripts upon T cell activation with single-cell and subcellular resolution.

**Fig. 1:**
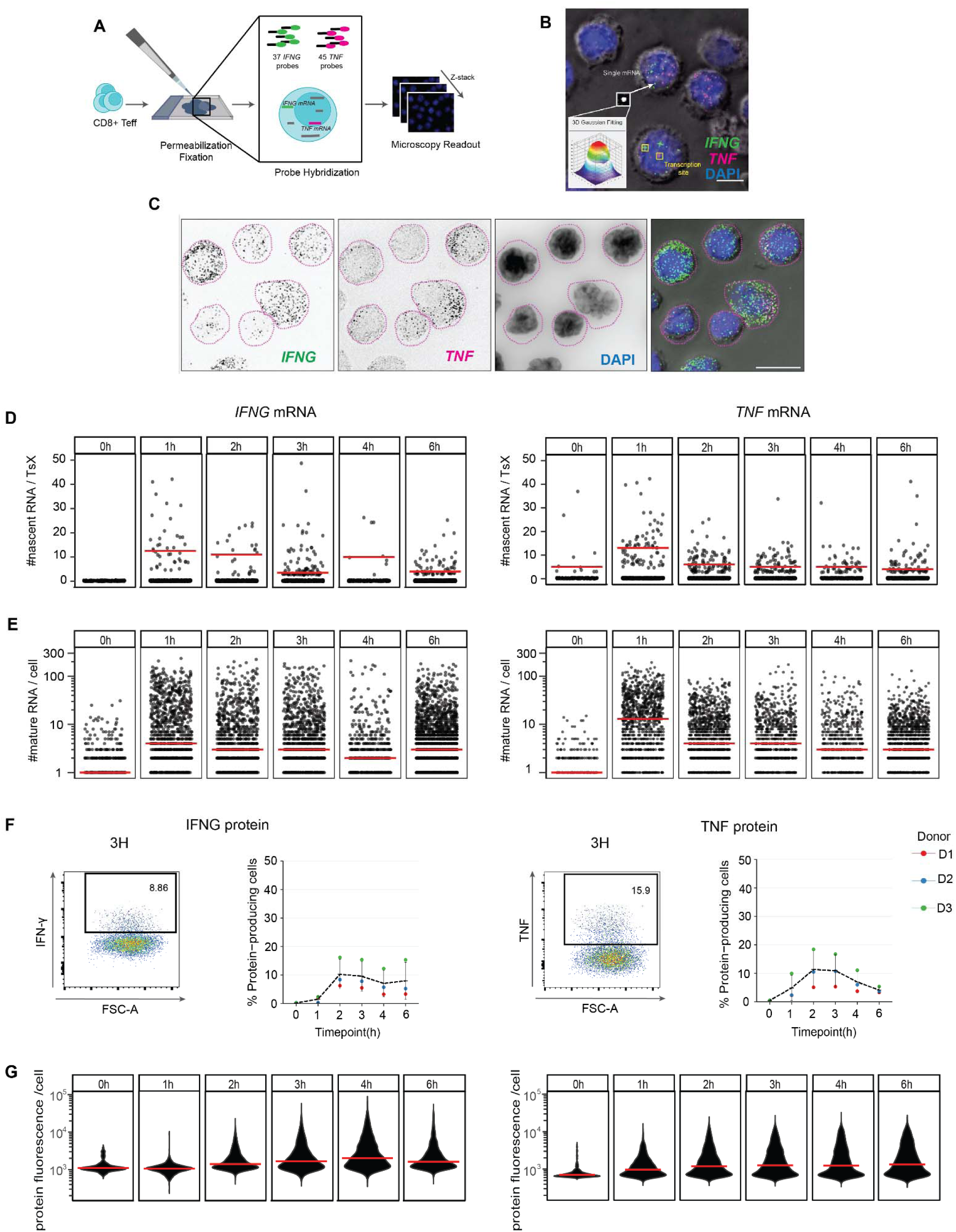
T-cell smFISH quantification of *IFNG* and *TNF* mRNA in CD8^+^ T eff cells. **A.** Schematic workflow of T-cell smFISH. **B.** Maximal projection of smFISH for *IFNG* mRNA (green), *TNF* mRNA (magenta), and DAPI (blue) merged to a single DIC section (grey). Yellow squares indicate *IFNG* and *TNF* transcription sites (TsX). Lower left inset: Gaussian signal of a single mature mRNA. Scale bar: 5 μm **C.** Maximal projection of smFISH for *IFNG* mRNA (first image, green), *TNF* mRNA (second image, magenta), DAPI (third image, blue) and all previous images merged on DIC. Dotted lines represent 2D cell outline. Scale bar: 10 μm **D.** Number of nascent *IFNG* (left) and *TNF* (right) RNA per TsX. Each dot depicts one transcription site. Data pooled from 3 donors. n ≥ 1000 cells/time point. Red bar: median. **E.** Number of mature *IFNG* (left) and *TNF* (right) mRNA in Teff cells expressing ≥1 mature mRNA. Each dot represents an individual cell. Data pooled from 3 donors. **F.** Cytokine expression measured by intracellular cytokine staining. Bref A was added for max. 2h. Left panels: representative flow cytometry plot. Right panels: n=3 donors, line= mean. **G.** Violin plot of protein fluorescence intensity per cell. Data pooled from 3 donors. Red bar: median.

### T-cell smFISH uncovers cytokine-specific transcription activity

Having established T-cell smFISH, we studied the cytokine (m)RNA expression changes during T cell activation with α-CD3/α-CD28. The key effector cytokines IFN-γ and TNF are produced early after T cell activation, yet with distinct kinetics (*28*, *29*). To uncover the underlying mechanism, we measured nascent RNA and the subcellular distribution of mature mRNA (**Fig. Suppl. 1D**), and we compared that to cytokine protein levels.

We first focused on nascent cytokine RNAs. Resting Teff cells (t=0 h) lacked active transcription sites (TsX) for *IFNG* (**Fig. 1D, Suppl. Fig. E**). At 1 h of activation, however, 2% Teff cells expressed nascent *IFNG* mRNA with a median of 12 nascent RNAs per TsX (**Fig. 1D; Suppl. Fig. 1E-1F**). At 3 h of activation, active TsX peaked with 8%, with a median of 3 nascent *IFNG* RNAs per TsX, and this level of nascent RNA was maintained up to 6 h of activation (**Fig. 1D, Suppl. Fig. 1F-1G**). Of note, even though at 1 h of activation we observed bi-allelic transcription in 45% of the *IFNG*-transcribing cells, at later time points monoallelic *IFNG* transcription was more prominent (**Suppl. Fig. 1E**).

In contrast to *IFNG,* nascent *TNF* RNA was already detected in 1% of resting Teff cells with a median of 5 nascent RNA molecules per TsX (**Fig. 1D, Suppl. Fig. 1E-1F**). At 1h of stimulation, nascent *TNF* peaked with 13 nascent RNA per TsX, which dropped and stabilized at 2h with 6 molecules on average per cell (**Fig. 1D, Suppl. Fig. 1E-1F**). Intriguingly, this drop did not result from fewer active TsX for *TNF*. Furthermore, bi-allelic transcription was observed in 56% of the TNF-transcribing cells until 3 h of activation, and only at later time points, mono-allelic transcription was more prevalent (**Suppl. Fig. 1E**). Thus, nascent *IFNG* and *TNF* RNA display different kinetics, with *IFNG* requiring T cell activation for *de novo* synthesis, and *TNF* transcription already being active in resting Teff cells. *IFNG* transcription relies more on mono- allelic transcription, whereas *TNF* transcription also employs bi-allelic synthesis during early T cell activation.

### Cytokine-specific kinetics of mRNA and protein expression during T cell activation

We next quantified mature *TNF* and *IFNG* mRNA. Even though nascent *IFNG* RNA was undetectable in resting Teff cells, 36% expressed low but detectable levels of mature *IFNG* mRNA, and 28% expressed mature *TNF* mRNA (each a median of 1 molecule/cell; **Fig.1E, Suppl. Fig. 1G**). At 1 h of stimulation, 77% and 74% of the T cells expressed mature *IFNG* and *TNF* mRNA, respectively (median of 4 and 13 molecules/cell, respectively; **Suppl. Fig. 1G-H**). Notably, a high inter-cellular variability was observed with 1-200 *IFNG* and *TNF* mRNA molecules/cell in all three donors, and at all measured time points (**Fig. 1E; Suppl. Fig. 1H)**. Yet, whereas a subset of Teff cells maintained high levels of mature *IFNG* mRNA throughout T cell activation (>100 molecules/cell; **Fig 1E**), the levels of mature *TNF* mRNA rapidly declined below 100 molecules/cell from 2 h of activation onwards (**Fig. 1E**).

In line with the literature (*28*, *29*), also the protein production of IFN-γ and TNF displayed intercellular and inter-donor heterogeneity (**Fig. 1F**). Yet, the overall response kinetics were comparable between donors. Following the expression patterns of mature mRNA, IFN-γ protein production peaked at 2 h of activation with a maximum of 10%, and it remained stable thereafter (**Fig. 1F**). Furthermore, also at protein levels a high inter-cellular variability was detected, which on population level peaked at 4h post-stimulation (**Fig. 1G, p<0.0001 compared to all other time points).** TNF production was more rapid: in line with higher *TNF* mRNA levels at 1 h of activation (**Fig. 1E, p<0.0001 compared to 0h**), TNF protein production peaked at 2-3 h of activation with 12% TNF-producing Teff cells, and the percentage rapidly declined thereafter (**Fig. 1F**). Nevertheless, the protein production per cell remained constant throughout (**Fig.1G**). Combined, these data indicate that the protein production kinetics of IFN-γ and TNF follow the cytokine mRNA kinetics.

### Dual cytokine mRNA-expressing Teff cells dominate the immune response

Dual cytokine producers are more potent effector T cells than single cytokine producers in response to infection and cancer (*45*, *46*). We therefore questioned how single (SP) and double positive (DP) cytokine mRNA expressors (**Movie S1**) correlate with the observed high variability of cytokine mRNA expression. When enumerating the number of nascent cytokine RNA, we found significantly higher numbers of nascent *IFNG* RNAs per TsX in DP compared to SP Teff cells (**Fig. 2A**). This was not the case for nascent *TNF* (**Fig. 2A**). In contrast, the percentage of active TsX was substantially higher for both cytokines in DP Teff cells, which was observed at all time points **(Suppl. Fig. 2A**). In addition, DP T cells displayed more bi-allelic transcription for both cytokines (**Suppl. Fig. 2A**). The higher transcriptional activity in DP T cells also resulted in significantly higher numbers of mature cytokine mRNA per cell (**Fig. 2B, C**). Upon T cell activation, DP T cells expressed substantially higher percentages and numbers of mature *IFNG* mRNA/cell at all time points measured and mature *TNF* mRNA at early T cell activation time points (1-3 h) compared to SP T cells (**Fig. 2B, C)**. Not only mRNA, but also the percentage of protein-producing Teff and protein production/cell was higher in DP cytokine mRNA producers (**Fig. 2D, Suppl. Fig. 2B**). In conclusion, Teff cells with active TsX for both *IFNG* and *TNF* contain more mature cytokine mRNA, and the higher mRNA levels originate from higher levels of biallelic transcription. Furthermore, DP cytokine producers dominate the cytokine production kinetics upon T cell activation and contribute to the observed high inter-cellular heterogeneity of cytokine production.

**Fig. 2:**
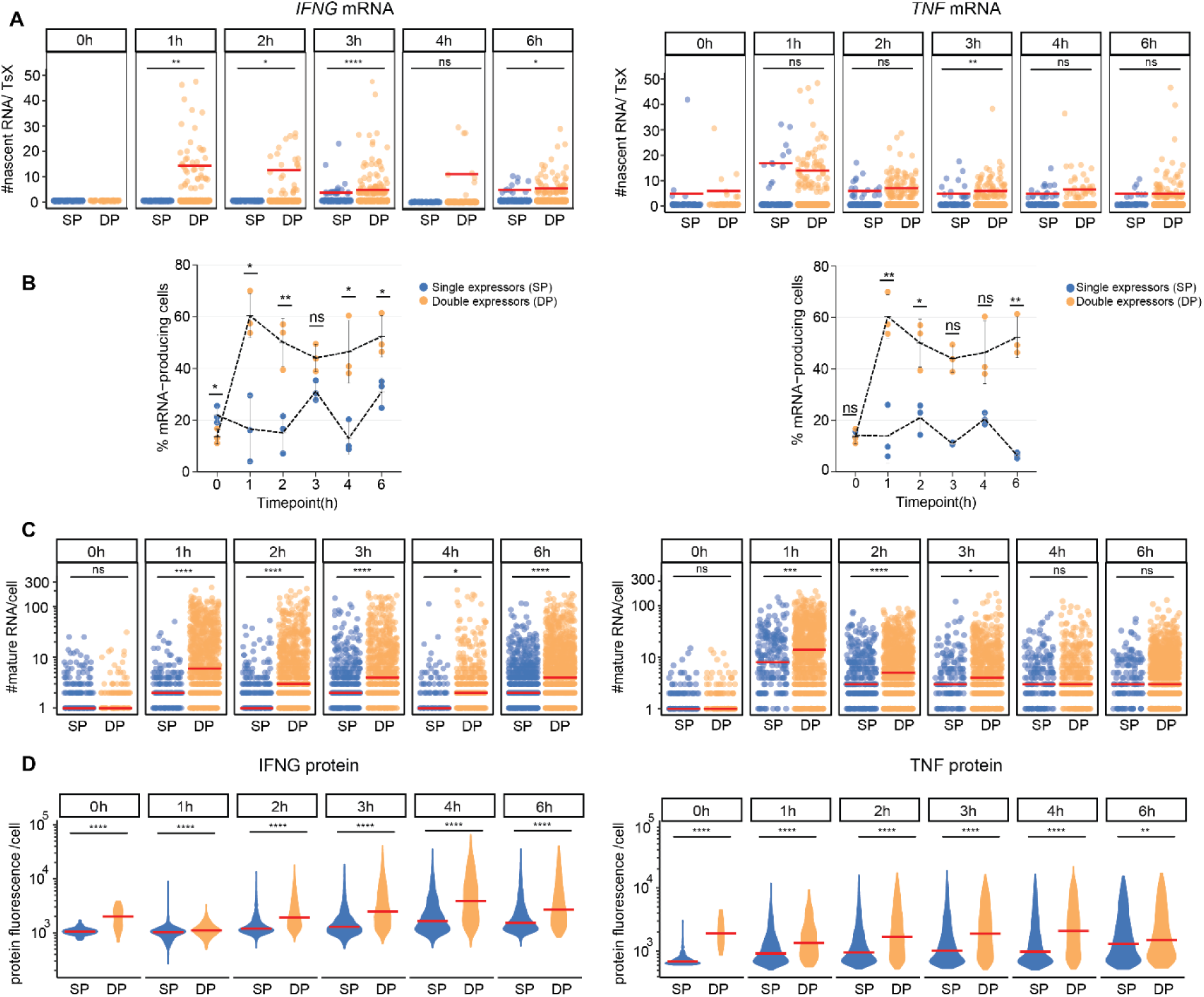
Cytokine mRNA and protein expression is dominated by double expressors. **A.** Number of nascent *IFNG* (left) and *TNF* (right) RNA per TsX in single positive (SP, blue) or double positive (DP, orange) expressors upon T cell activation. Each dot represents an individual cell, data pooled from 3 donors. Red bar: median. **B.** Percentage of Teff cells expressing ≥1 mature *IFNG* (left) and *TNF* (right) mRNA in single and double positive mRNA expressors. n=3 donors, line: mean. **C.** Number of mature *IFNG* (left panel) and *TNF* (right panel) mRNA SP and DP expressors containing ≥1 mRNA. Each dot represents an individual cell. Data pooled from 3 donors. Red bar: median. **D.** Violin plot of protein fluorescence intensity of cytokine production in SP and DP producers per cell. Data pooled from 3 donors. Red bar: median. *p≤0.05, **p≤0.01, ***p≤0.001, ****p≤0.0001 ns: non-significant. Kruskal-Wallis non-parametric test, and post-hoc Tukey HSD test for comparing time points.

### Differential subcellular distribution of *IFNG* and *TNF* mRNA

Mature cytokine mRNA can only serve as template for translation when located in the cytoplasm. Therefore, we next investigated the subcellular distribution of *IFNG* and *TNF* mRNA during activation with T-cell smFISH (**Suppl. Fig. 3A**). In resting Teff cells, the vast majority of *IFNG* mRNA and *TNF* mRNAs were present in the nucleus (**Fig. 3A, Suppl. Fig. 3A - top panel**). However, already 1 h of T cell activation rapidly shifted the distribution of mature *IFNG* mRNA to the cytoplasm (**Fig. 3A)**, indicating rapid translocation after T cell activation. From 2 h of activation onwards, the *IFNG* mRNA was equally distributed between nucleus and cytoplasm (**Fig. 3A**). *TNF* mRNA was more prevalent in the nucleus throughout the activation phase, which was with 75% nuclear mRNA most apparent at the peak of its expression, i.e. at 1 h of T cell activation (**Fig. 3A, Suppl. Fig. 3A**). Thus, mature *IFNG* and *TNF* mRNAs differentially distribute between nucleus and cytoplasm in activated Teff cells.

**Fig. 3:**
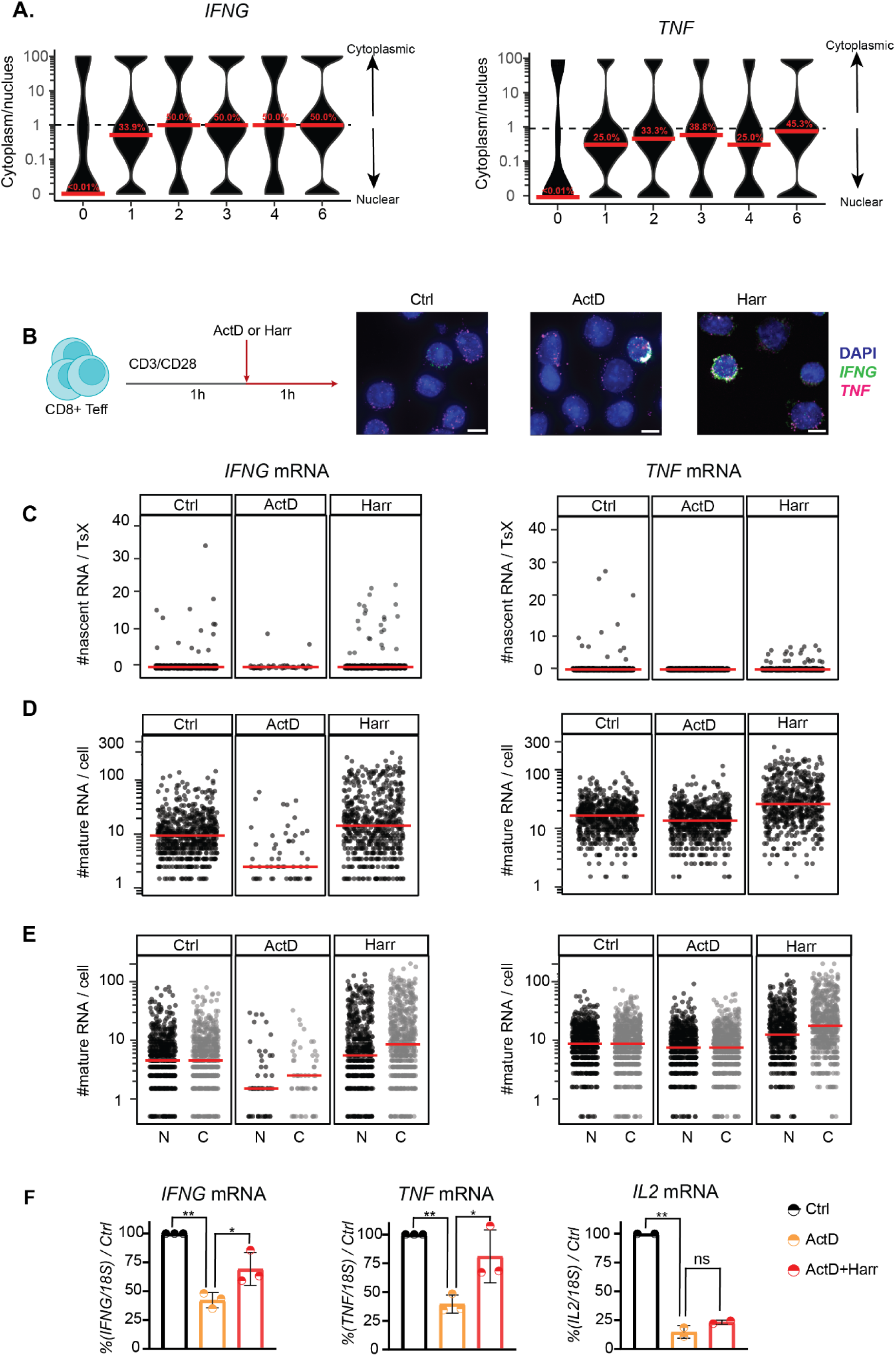
*IFNG* and *TNF* mRNA localization and translation control. **A.** Ratio of cytoplasmic mRNA versus nuclear mRNA per cell of Teff cells expressing ≥1 mature mRNA. Data represented as pseudo-log10 transformed. Dotted line represents equal distribution in nucleus and cytoplasm. n ≥ 500 cells/time point, pooled from 3 donors. Red bar: median. Percentage in red: median of percentage of mRNA in the cytoplasm. **B.** Left: Experimental setup of transcription/translation block in Teff cells. Right: MAX projections of T-cell smFISH. Scale bar; 5μm. **C.** Number of nascent *IFNG* (left) and *TNF* (right) RNA per TsX. Each dot represents an individual cell. n ≥ 1000 cells/condition, one donor. Red bar: median. **D.** Mature cytokine mRNA of Teff cells expressing ≥1 mature mRNA. **E.** Number of nuclear (N, black) and cytoplasmic (C, grey) cytokine mRNA. **F.** Cytokine mRNA expression measured by qRT-PCR upon treatment with ActD, or ActD for 20min then combined with Harr, normalized to control (Ctrl). n=3 donors, mean ± SD, p≤0.05, **p≤0.01, ***p≤0.001, ****p≤0.0001 ns: non- significant. Two-tailed ratio unpaired student t-test.

### Translation-dependent decay controls cytokine mRNA levels upon T cell activation

The unexpectedly high levels of mature cytokine mRNA in the nucleus prompted us to question how *de novo* transcription and translation contributed to the subcellular distribution. We therefore activated Teff cells for 2h, and in the 2^nd^ hour of activation, we blocked either *de novo* transcription with Actinomycin D (ActD), or we blocked translation with Harringtonine (Harr) (**Fig. 3B**). As expected, Harr treatment reduced both the overall translation as defined by puromycin incorporation, and the cytokine production (**Suppl. Fig. 3B, C**). Likewise, T-cell smFISH analysis confirmed that ActD almost completely blocked the presence of nascent RNA (**Fig. 3C**). ActD treatment also substantially reduced the number of mature *IFNG* mRNA molecules/cell (**Fig. 3D**). Intriguingly, the mRNA expression of mature *TNF* was much less affected by ActD treatment (**Fig. 3D)**. Because *TNF* mRNA was not found to be stabilized upon T cell activation (*6*), the data suggest that the production boost of *TNF* mRNA we observed during the first hour of T cell activation (**Fig. 1E**) suffices to maintain high levels of mature *TNF* mRNA expression during the 2^nd^ hour of activation.

Notably, blocking translation with Harr increased the numbers of mature cytokine mRNA/cell compared to control (**Fig. 3D, p<0.0001).** T-cell smFISH uncovered that Harr treatment primarily increased the cytoplasmic *IFNG* mRNA and *TNF* mRNA (**Fig. 3E, p<0.0001**), suggesting that the effect of Harringtonine on cytokine mRNA expression resulted from blocking translation. Previous studies reported that the cytoplasmic RNA turnover can be influenced by translation (*47*, *48*) . To determine whether the observed cytoplasmic accumulation of cytokine mRNA in T cells upon Harr treatment resulted from translation-dependent mRNA decay, we first blocked transcription and subsequently translation and measured cytokine mRNA by qRT-PCR. ActD treatment alone reduced the overall cytokine mRNA levels by 60% compared to control (**Fig. 3F, p<0.01**), however, this reduction had little effect on the protein output (**Suppl. Fig. 3D, E**). In contrast, ActD/Harr treatment combined substantially reduced the protein production (**Suppl. Fig. 3D, E)**. Yet, the expression levels for *IFNG* and *TNF* mRNA increased by 30% in ActD/Harr-treated Teff cells compared to Act treatment alone (p<0.05), suggesting that *IFNG* and *TNF* mRNA availability for translation is regulated by translation. Of note, this increase was not observed for *IL2* mRNA, which suggests transcript specificity for this process (**Fig. 3F)**. Combined, the T-cell smFISH pipeline uncovered - with single-cell resolution - that translation regulates the cytoplasmic *IFNG* and *TNF* mRNA levels. Furthermore, *IFNG* and *TNF* mRNA - but not *IL2* mRNA- are subject to translation-dependent decay.

### HuR differentially controls *IFNG* and *TNF* mRNA expression and cytokine production

Having established the cytokine mRNA expression kinetics and distribution in Teff cells, we questioned how RNA-binding proteins such as HuR regulate the fate of cytokine mRNA. We previously showed that HuR promotes cytokine production during the early phase of T cell activation, i.e. at 1-2h, but not at later time points (*6*), Thus, HuR activity supports the rapid response rate of effector T cells. Its mode of action is, however, unresolved. HuR protein expression increased in Teff cells during the 2 h of activation (**Suppl. Fig. 4 A)**. However, RNA-immunoprecipitation (RIP)-qRT-PCR revealed that HuR only interacted with *IFNG* and *TNF* mRNA at 1 h post stimulation, and not at 2h post-stimulation (**Suppl. Fig. 4 B**). To study its mode of action on cytokine mRNA, we depleted HuR from Teff cells by CRISPR-Cas9 gene editing (**Suppl. Fig. 4C)**. As previously reported (*6*), HuR-KO Teff produced less IFN-γ and TNF protein during the first 2h of activation, both in terms of percentage and of protein production/cell **(Fig. 4A, B)**. Notably, T-cell smFISH analysis uncovered opposite effects on mRNA levels (**Fig. 4C, D)**. Resting HuR-KO Teff cells already contained a higher percentage of *IFNG* mRNA expressing cells than control-treated T cells (**Suppl. Fig. 4D).** The median mRNA molecules/cell increased from 3 to 10 for *IFNG*, and from 5 to 9 for *TNF* mRNA (**Fig. 4D)**. At 1h of stimulation, the percentage of mRNA-expressing Teff cells and number of cytokine molecules/cell was also higher in HuR-KO (**Fig. 4D, Suppl. Fig. 4D).** Only at 2 h activation, the numbers of cytokine mRNAs/cell dropped below those of control Teff cells (**Fig. 4D)**.

**Fig. 4:**
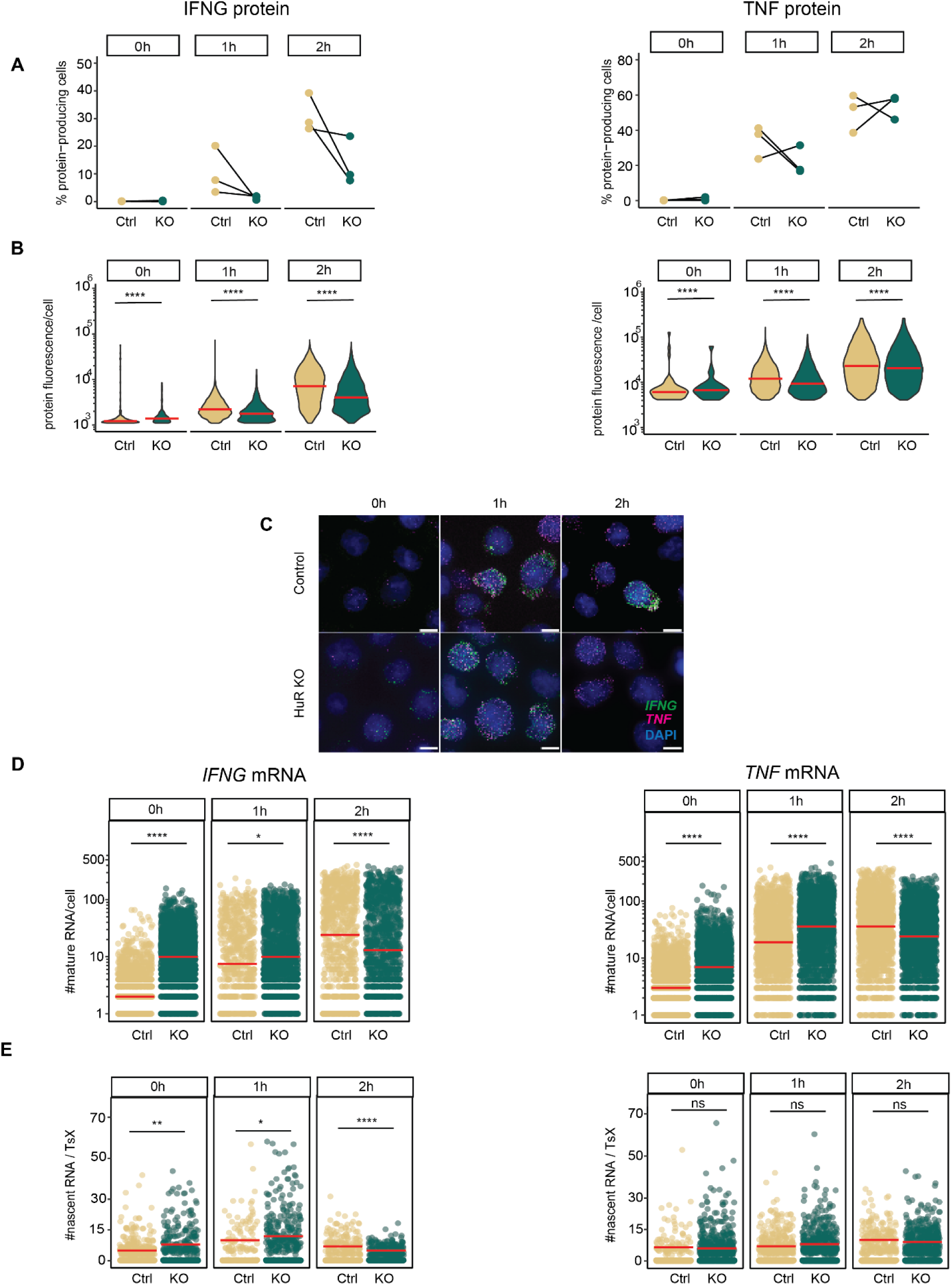
IFNG and TNF mRNA and protein expression HuR-KO Teff cells. **A.** Percentage of cytokine-producing Ctrl (C) and HuR-KO Teff cells (KO). BrefA was added at 0h. n=3 donors. ns: non-significant. Kruskal-Wallis non-parametric test, and post-hoc Tukey HSD test. **B.** Violin plot of protein fluorescence intensity per cell, from 3 pooled donors. Red bar: median **C.** Control (top panel) and HuR KO (bottom panel) Teff cells were activated with α- CD3/α-CD28. MAX projection of T-cell smFISH. Scale bar: 5μm. **D.** Number of mature cytokine mRNA in Teff cells expressing ≥1 mature mRNA. n ≥ 1500 cells/time point, from 3 pooled donors. Red bar: median. **E.** Number of nascent cytokine RNA per TsX. Red bar: median. *p≤0.05, **p≤0.01, ***p≤0.001, ****p≤0.0001 ns: non-significant. Kruskal-Wallis non- parametric test, and post-hoc Tukey HSD test for comparing time points.

To determine whether the higher cytokine mRNA levels observed in HuR-KO Teff cells resulted from increased transcription, we measured the number of nascent RNA. Resting HuR-KO Teff cells had on average 8 nascent *IFNG* RNA/cell compared to 5 nascent *IFNG* RNA/cell in control T cells, and this slight increase was lost by 2h post activation (**Fig. 4E**). In contrast, nascent *TNF* RNA remained unaffected by HuR depletion at all time points (**Fig. 4E)**. Thus, despite that HuR-KO T cells produce less cytokine during early activation, mature mRNAs accumulate, and transcription may only partly contribute to this accumulation.

### HuR deletion affects the cellular distribution of *IFNG* and *TNF* mRNA

Because higher cytokine mRNA levels in HuR-KO cells did not result in higher but lower protein production, we questioned whether HuR depletion affected the subcellular mRNA distribution. T- cell smFISH analysis uncovered that resting HuR-KO Teff cells expressed significantly more cytoplasmic *IFNG* and *TNF* mRNA molecules than control Teff cells (**Fig. 5A**). HuR-KO Teff cells also contained slightly more nuclear *TNF* mRNA (**Fig. 5A**). Upon 1h activation, both cytokine mRNAs were slightly increased in both subcellular compartments, which was reversed at 2 h of T cell activation (**Fig. 5A**). When we determined the relative distribution of cytokine mRNA within each individual Teff cell, we found that resting HuR-KO T cells contain more *IFNG* mRNA in the cytoplasm, which drops at 1h but is reverted at 2h post-stimulation (**Fig. 5B)**. In contrast, *TNF* mRNA is slightly more nuclear at 0h and 1h of activation in HuR-KO cells compared to control cells, and this effect is lost at 2h of activation, at a time point when HuR also loses its interaction with the cytokine mRNAs (**Fig. 5B, Suppl. Fig. 4B**). In conclusion, HuR deletion influences the subcellular localization of cytokine mRNAs, and it does so in a time- dependent manner.

**Fig. 5:**
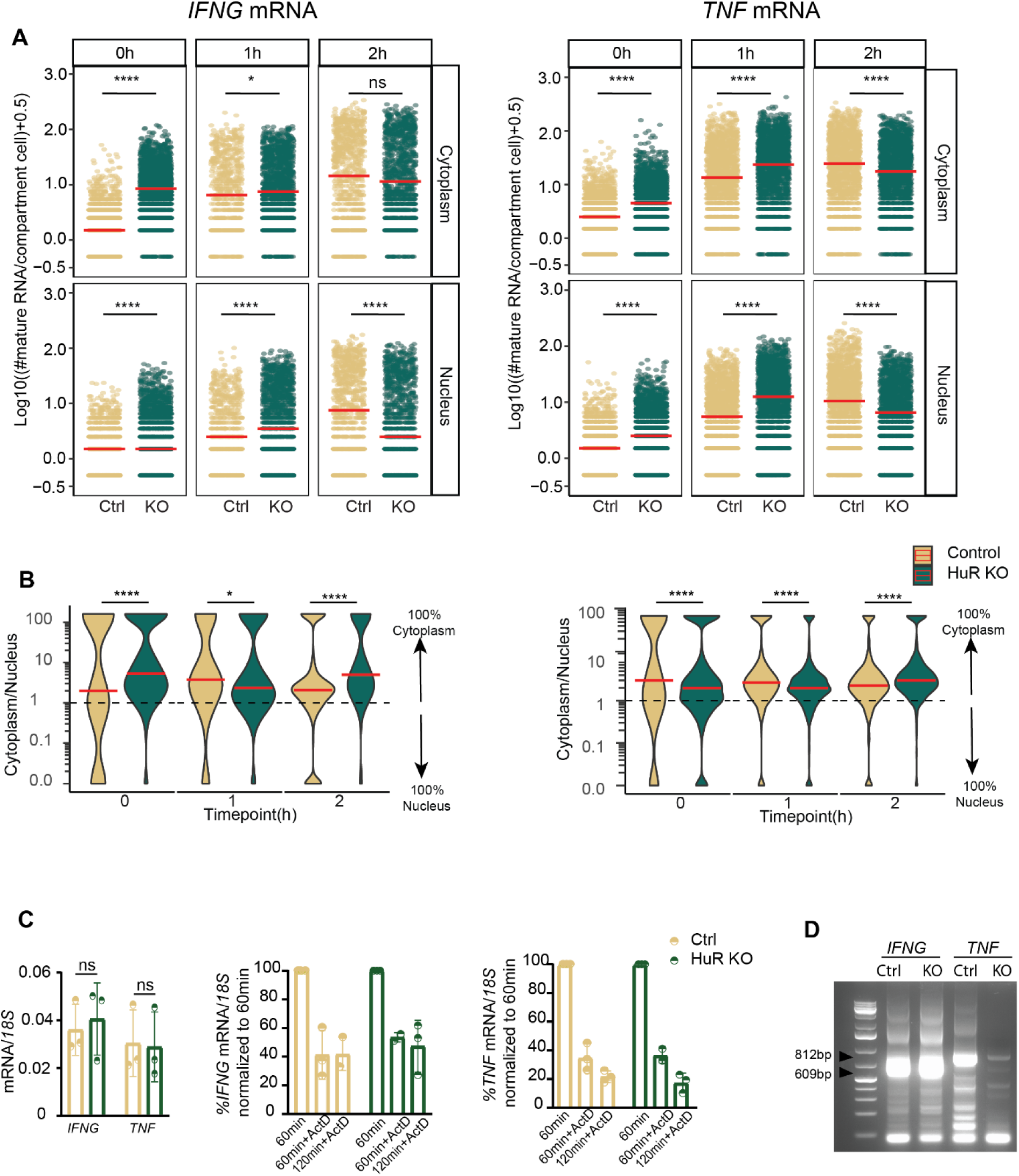
*IFNG* and *TNF* mRNA localization and PTR in HuR-del Teff cells. **A.** Number of cytoplasmic (top) and nuclear (bottom) cytokine mRNAs in control (Ctrl) or HuR- KO (KO) Teff cells that express ≥1 mature mRNA per cell. Data represented as pseudo-log10 transformed, with a coefficient of 0.5 added. Each dot represents an individual cell, data pooled from 3 donors. Red bar: median. **B.** Ratio of cytoplasmic mRNA versus nuclear mRNA in Ctrl or HuR-KO Teff cells expressing ≥1 mature mRNA per cell. Data represented as pseudo-log10 transformed. Data from 3 pooled donors. Red bar: median. For Fig. A-B *p≤0.05, **p≤0.01, ***p≤0.001, ****p≤0.0001 ns: non-significant. Kruskal-Wallis non-parametric test, and post-hoc Tukey HSD test for comparing time points. **C.** Left: *IFNG* and *TNF* mRNA expression determined by qRT-PCR in Ctrl or HuR-KO Teff cells activated for 1h with α-CD3/α-CD28. Right *IFNG* and *TNF* mRNA levels in Teff upon ActD treatment. Cytokine mRNA expression at 1h of T cell activation is considered 100%. n=3 donors, mean ± SD, Data non-significant. Two- tailed ratio paired t-teest **D.** Measurement of poly(A) length of cytokine mRNA using RL-PAT in control (Ctrl) or HuR-KO (KO) Teff cells activated for 1h with α-CD3/α-CD28 using the RL- PAT assay. Arrows indicate *IFNG* 3’UTR full-length: 812bp and *TNF* 3’UTR full-length: 609bp.

### HuR regulates the polyadenylation of *TNF* mRNA

Lastly, we aimed to decipher how HuR regulates cytokine RNA. HuR can stabilize its target mRNAs (*49*, *50*). However, overall mRNA levels at 1h of activation were not altered (**Fig. 5C**). Furthermore, measuring mRNA decay in T cells activated for 1h, and then treated for an additional 1-2h with ActD showed no differences in RNA decay rates between HuR-KO and control Teff cells (**Fig. 5C**). Thus, HuR does not appear to regulate cytokine mRNA stability.

HuR can also act as a splicing factor (*19*, *51*). Yet, irrespective of the nuclear *IFNG* mRNA accumulation (**Fig. 5B**), no overt differences of intron-exon junctions were found between HuR- KO and control Teff cells (**Suppl. Fig. 5A)**. Additionally, even though splicing is considered a key event for *TNF* (*52*, *53*), HuR deletion has only limited- if any- effect on *TNF* splicing (**Suppl. Fig. 5A),** indicating that HuR is not essential for *IFNG* and *TNF* RNA splicing in T cells. Lastly, HuR can regulate polyadenylation of its target mRNA (*18*, *21*). For *IFNG* mRNA, we found no changes in the poly(A) tail profile in HuR KO cells at 1h of T cell activation (**Fig 5D, Suppl. Fig 5B**). In sharp contrast, HuR deficiency results in changes of the poly-adenylation profile of *TNF* mRNA, as defined by the RNA ligation-mediated poly(A) test (RL-PAT), which measures the poly(A) tail length of mRNAsA novel method for poly(A) fractionation reveals a large population of mRNAs with a short poly(A) tail in mammalian cells (*54*). Thus, HuR deficiency impairs the canonical poly(A) tail. (**Fig 5D, Suppl. Fig 5B**). Thus, HuR employs cytokine-specific regulatory mechanisms to control the expression of mRNA, and ultimately the protein expression of these two key pro-inflammatory cytokines.

## Discussion

In this study, we present the T-cell smFISH pipeline to simultaneously quantify nascent and mature cytokine mRNAs and their subcellular distribution. Importantly, the three-dimensional analysis of RNA expression with single-molecule and single-cell resolution provided new insights in the regulation of cytokine production and the activity of RNA binding proteins in T cells, as we showcased for HuR.

Even though most activated Teff cells express cytokine mRNAs, they display a two order of magnitude difference in mRNA and protein expression. Within these heterogenous T cell responses, cytokine production upon T cell activation primarily originates from Teff cells that transcribe both cytokine mRNAs simultaneously, suggesting that this superior cytokine production by DP cells is cell-intrinsic. Even though the underlying mechanism is yet to be uncovered, it is tempting to speculate that DP cells produce more cytokines because of 1) an open chromatin structure at the cytokine loci or their respective regulators, 2) high expression of cytokine-driving transcription factors, or 3) more responsive signaling cascades in response to T cell activation. Future T-cell smFISH analyses may support the quest to uncover the underlying mechanisms.

T-cell smFISH uncovered that *IFNG* and *TNF* mRNA accumulates in the nucleus, suggesting that nuclear retention regulates the availability of cytokine mRNAs for translation, as previously shown in other cellular system (*55*). In addition, nuclear retention of *TNF* mRNA may support its efficient splicing as previously suggested (*52*, *53*). We therefore hypothesize that the subcellular localization of cytokine mRNA dictates the amount of cytokine production. Furthermore, we here uncovered that translation-dependent decay (*56*) contributes to the lower cytoplasmic mRNA levels we measured, a feature that appeared transcript-specific, as *IL2* mRNA was resistant.

T-cell smFISH also provides novel insights in the mode of action of RBPs. HuR deficiency resulted in increased levels of *IFNG* and *TNF* mRNA, yet with lower protein production. Of note, smFISH probes can detect mature and pre-spliced RNA (*30*, *41*). However, splicing was not affected in HuR KO T cells. Intriguingly, T-cell smFISH revealed cytokine-specific regulation. HuR deficiency slightly altered the transcription of *IFNG* mRNA, but not that of *TNF* mRNA. In turn, HuR is required for the polyadenylation of *TNF*, but not of *IFNG*. Polyadenylation can modulate mRNA stability (*57*), but mRNA stability was not altered for *TNF*. HuR can also promote alternative poly-adenylation and compete with the PolyA cleavage factor (*18*, *58*). Whether these changes in *TNF* polyadenylation are directly linked to the observed impaired translation of TNF is, however, yet to be determined. Conversely, *IFNG* mRNA does not depend on HuR for splicing or polyadenylation. Therefore, HuR may regulate the nuclear export of *IFNG* mRNA during early T cell activation, as previously described for other target mRNAs (*59*, *60*). Irrespective of its exact mode of action on *IFNG* mRNA, our data highlight that HuR’s activity is target-specific, as was previously reported for other RBPs (*24*, *61*).

In summary, T-cell smFISH is a sensitive and quantitative tool with single-cell, single- molecule resolution for deciphering post-transcriptional regulation in T cells. We found that differential mechanisms govern *IFNG* and *TNF* mRNA kinetics and showcased the contribution of HuR in regulating the cytokine production of T cells in a transcript-specific manner. We propose that T- cell smFISH is a powerful tool to uncover molecular mechanisms underlying dysregulated cytokine production, such as impaired production upon chronic stimulation (*62*) or excessive production in immunopathology (*15*, *63*, *64*). The custom-made T-cell smFISH pipeline we present here could support further studies on RNA expression and RBP-mediated regulation that define T cell function.

## Material and Methods

### T cell activation and cell culture

Human T cells from anonymized healthy donors were used in accordance with the Declaration of Helsinki (Seventh Revision, 2013) after written informed consent (Sanquin). Peripheral blood mononuclear cells (PBMCs) were isolated by Lymphoprep density gradient separation (Stemcell Technologies). To generate Teff cells, CD8^+^ T cells were enriched from cryopreserved, defrosted PBMCs with the CD8^+^ T cell isolation kit (Miltenyi Biotec) with a purity of >85%. T cells were activated for 72 h with 1 μg/mL plate-bound α-CD3 (HIT3a) and 1 μg/mL soluble α-CD28 (CD28.2; Biolegend), as previously described (*6*). Cells were cultured at 37°C, 5% CO2 in culture medium (IMDM, GIBCO, Thermo Fisher Scientific, supplemented with 10% fetal bovine serum (FBS), 100 U/mL penicillin, 100 μg/mL streptomycin, 2 mM L-glutamine). Cells were harvested and cultured at a density of 1,5×10^6^/mL for 7 days in standing T25/75 tissue culture flasks (Thermo Scientific) in culture medium supplemented with 100 IU/mL recombinant human (rh) IL-2 (Proleukin). Medium was refreshed every 2 days. Upon nucleofection, T cells were cultured in T cell mixed media (Miltenyi Biotec) supplemented with 5% heat-inactivated human serum, 5% FBS, 100 U/mL Penicillin, 100 μg/mL streptomycin, 2 mM L-glutamine, 100 IU/mL rhIL-2.

### Gene-editing of primary human CD8^+^ T cells

Cas9 RNP production and nucleofection was performed as previously described (*6*). sgRNA targeting HuR (5’- TGTGAACTACGTGACCGCGA-3’ (*6*)) was dissolved in Nuclease Free Duplex buffer (Integrated DNA Technologies, IDT). Non-targeting negative control crRNA #1 (IDT) was mixed with tracrRNA at equimolar ratios (100uM) in nuclease-free PCR tubes and denatured at 95°C for 5 min. Nucleic acids were cooled down to RT prior to mixing and incubating for at least 10 min at RT with 30 μg Alt-R™ S.p. Cas9 Nuclease V3 (IDT). 3×10^6^ activated T cells (72 h) were nucleofected with the generated Cas9 ribonuclear proteins (RNPs) in 20μl P2 buffer (Lonza) in 16-well strips using program EH100 in a 4D Nucleofector X unit (Lonza). HuR depletion was confirmed on day 5 after nucleofection by flow cytometry, as described below.

### T cell activation

T eff cells were stimulated with 1μg/ml soluble α-CD3 (Pelicluster CD3, Sanquin) and 1 μg/mL soluble α-CD28 in culture media for indicated time points. For intracellular cytokine staining measurements, 1 μg/mL brefeldin A (BD Bioscience) was added during the last two hours of activation. For translation efficiency, T cells were incubated with Puromycin dihydrochloride (Sigma) for 10 min at 37°C. For mRNA stability measurements, T cells were activated for 1h with α-CD3/ α-CD28 and then treated for 1h with 5 mg/mL actinomycin D (ActD) (Sigma-Aldrich) and/or with 5μg/mL of Harringtonine (Abcam).

### smFISH probe design

Probes for *IFN* and *TNF* were previously described (*28*), and sequences are provided in **Table S1**. Briefly, probes were designed using Stellaris Probe Designer tool (from LGC Biosearch Technologies). Subsequent probe blasting resulted in removal of a couple of probes that showed predicted high affinity for secondary target genes. This resulted in 37 and 45 probes for *IFNG* and *TNF*, respectively. Because we observed autofluorescent granule structures in activated T cells in the FITC and Cy3 channels, CALfluorRed 610 and Quasar 670 were used as conjugates for IFNG and TNF probes, respectively (**Suppl. Material and Methods)**. Probes were purchased from LGC BioSearch Technologies.

### Two-color single molecule fluorescent in situ hybridization (smFISH)

Two-color smFISH was performed as described in with the following adjustments (*42*, *65*). Optimization of T cell attachment, cell amount and fluorophore selection are described in **Suppl. Materials and Methods.** RNase-free reagents were used throughout. Round 16-mm coverslips (Fisherbrand Borosilicate Glass Circle Coverslip) were cleaned with 0.5M HCl, washed twice with Milli-Q water, and coated with Alcian-blue (Alcian blue in 3% acetic acid, Sigma) for 30 min at RT. Upon extensive washes with Milli-Q water, 1×10^6^ Teff cells were seeded in 60 μl ice-cold 1xPBS per coverslip to prevent autolysis and stop any ongoing cellular process. T cells were incubated for 20 min at 37°C to enhance T cell attachment. Coverslips were gently washed with ice-cold PBS to remove non-attached cells. Teff cells were fixed with 4% paraformaldehyde (32% solution, EM grade; Electron Microscopy Science #15714) for 10 min at RT in the dark.

Cells were washed twice with quenching buffer (3mM MgCl_2_, 0.1M glycin in 1xPBS) for 5 min at RT. Coverslips were gently covered with 70% ice-cold ethanol and stored at -20°C until further use, both to decrease autofluorescence, and to preserve RNA and cellular structure. For hybridization, coverslips were rehydrated by washing twice with 2xSSC for 5 min prior to incubation with pre-hybridization buffer (2xSSC, 10% deionized formamide, 1:1000 RNase-inhibitor in ultrapure distilled water) for 50 min at RT. IFNγ-CF610 and TNF-Quasar 670 probes were freshly diluted to a final concentration of 125nM in hybridization solution A. Hybridization solution A contains 20% formamide to denature single-strand DNA, and was supplemented with 10μg/ml yeast tRNA (Invitrogen) and 10μg/ml salmon sperm DNA (Ultrapure salmon sperm DNA solution, Invitrogen) used as nucleic acid competitors to saturate nonspecific probe binding. Hybridization solution A was denatured for 5 min at 95°C. Hybridization solution A was added to 1:1 ration with Hybridization solution B. Hybridization solution B includes reagent that are temperature sensitive. Specifically, we use 10% Dextran Sodium Sulfate to enhance probe hybridization, 10mg/ml BSA to reduce background signal, 1:1000 murine RNase inhibitor to preserve RNA and 2xSSC in ultrapure water). Hybridization was performed for 3 h at 37°C in the dark. Coverslips were washed twice with pre-hybridization buffer for 15 min at 37°C, followed by two washes with 2xSSC for 10 min at RT. After a last wash in 1xSSC for 5 min at RT, cells were incubated with 1μg/ml DAPI (Thermo Scientific) for 5 min in the dark at RT. Coverslips were washed twice at RT with 1xPBS and mounted on glass with Prolong Diamond Antifade mountant (Invitrogen).

### T cell smFISH image acquisition and analysis

Images were acquired as previously described (*43*). Briefly, image acquisition was performed using an Olympus BX-63 epifluorescence microscope equipped with Ultrasonic stage and UPLSAPO 100X1.4NA oil immersion objective (Olympus). Lumencore SOLA FISH light source, Hamamatsu ORCAFusion sCMOS camera (6.5 μm pixel size) mounted using U-CMT C-Mount Adapter, and zero-pixel shift filter sets: F36-500 DAPI HC Brightline Bandpass Filter, F36-502, FITC HC BrightLine Filter, F36-542 Cy3 HC BrightLine Filter, AHF-LED-FISH-R Filter for Cy3.5 and F36-523 Cy5 HCBrightLine Filter. Z- section of 200nm intervals over an optical range of 12 μm. Per each coverlip, 12 positions of 61 Z-stack were collected to obtain measurements from at least 500 cells per each donor per time point. The CellSens software (Olympus) is used for instrument control and image acquisition.

sm-FISH images were analysed with FISH-quant Matlab (*41*). MAX projections of fluorescent background were used to obtain 2D-cellular outline using the fq-segmentation pipeline from FISH-quant (https://fish-quant.github.io/). After background subtraction, the Transcript Site Outline tool (FISH-quant) was used to identify transcription sites as high-intensity signals (x1.5 intensity of averaged mRNA) colocalizing with DAPI staining. Mature RNA quantification was performed by fitting RNA spots to a three-dimensional (3D) Gaussian. The intensity and width of the 3D Gaussian tool of FISH-quant were thresholded to exclude nonspecific signals. To quantify the number of nascent RNA per each Transcription sites, the average intensity of all mRNAs was used as a reference. Data were post-processed using Filtering.Rmd script (https://github.com/nikolinasostaric/T-cell_smFISH). With the 2D-outline output from FISH- quant, cells in division (with two DAPI stained nuclei), cells on the edge of the coverslips, and cells with miscalled nuclei were excluded from further analysis (∼20% of the cells/experiment).

Subcellular localization of RNA was defined with the 3D spot localization pipeline. 3D segmentation of nuclei was performed with CellPose v2.2 (https://www.cellpose.org/) (*44*). DAPI staining over *z-*stacks was used for the nuclear outline. The following settings were employed: cytoplasm model 2.0 (cyto2), 3D setting stitch_threshold > 0, and nuclei diameter were calculated as the average diameter of images per each time point of activation. Nuclear mask coordinates and coordinates (*x,y,z*) per each RNA spot were imported into Spot_localization.ipynb script. *x,y* and *z* coordinates were used to define the colocalization of each mRNA with nuclear mask (per respective image). mRNA spots with coordinates not matching the nuclear mask were considered cytoplasmic mRNA.

### Quantitative (RT-qPCR) and reverse transcription- PCR (RT-PCR) analysis

Total RNA was extracted from T cells using the Quick-RNA Miniprep Kit (Zymo Research, R1055) according to the manufacturer’s protocol. cDNA was generated with Maxima First Strand cDNA Synthesis Kit (ThermoFisher Scientific, K1642). To study cytokine mRNA expression, RT-qPCR was performed with duplicate reactions using Power SYBR Green (Applied Biosystems, 4367659) on a StepOne Plus (Applied Biosystems). Ct values were normalized to 18S levels. *IFNG*, *TNF* and *IL2* primers used were previously described . For splicing RT-PCR, 200 ng cDNA was amplified with GoTaq G2 Flexi polymerase (Promega, M7805) with human *IFNG* and *TNF* exon- and intron-specific primers, designed by using the Primer3Plus (*66*) (**Table S2**) with the following protocol: 95°C for 1 min, 30 cycles of (30 sec at 95°C, 30 sec at 50°C, 2.5 min at 72°C), followed by 5 min at 72°C. PCR products were run on a 1-1.2% agarose gel. Quantification was performed with Analyze-Gel of Fiji version 2.15.1.

### RNA ligation-mediated poly(A) test (RL-PAT)

RL-PAT was performed as described (*54*). The 5’ to 5’ adenylated and 3’ blocked ‘PAT anchor’ oligo was ligated to the 3’ end of total RNA overnight at 16°C with RNA ligase 2, truncated KQ (NEB, M0373). To generate cDNA, ligated RNA was reverse transcribed with SuperScript III Reverse Transcriptase (Invitrogen, 18080044) with the ‘PAT-R1’ oligo (complementary to ‘PAT anchor’). cDNA was then amplified with GoTaq G2 Flexi Polymerase (Promega, M7805), using a forward primer annealing to the 3’ UTR of the mRNA of interest (**Table S3**) and PAT-R1 as the reverse primer. All mRNA specific PAT primers were validated by performing PAT on mRNA deadenylated with oligo-d(T) and RNAse H.

### RNA immunoprecipitation and immunoblotting

RNA immunoprecipitation was performed as previously described (*6*). Briefly, cytoplasmic lysates of 20×10^6^ human CD8^+^ Teff cells were prepared with lysis buffer (10 mM HEPES, pH 7.0, 100mMKCl, 5mMMgCl2, 0.5% NP40) freshly supplemented with 1mMDTT, 100 U/ml RNase OUT (both Invitrogen), 0.4Mm Ribonucleoside Vanadyl Complex (NEB) and 1% EDTA-free protease/phosphatase inhibitor cocktail (Thermo Scientific). Protein G Dynabeads (Invitrogen) were prepared according to manufacturer’s protocol. The lysate was immunoprecipitated for 4h at 4°C with 10 mg mouse monoclonal a-HuR (3A2, Santa Cruz Biotechnology) or a mouse IgG1 kappa isotype control (P3.6.2.8.1, eBioscience). RNA was extracted from beads with Trizol, and mRNA expression was measured by RT-PCR. The specificity of the RNA-IP was validated by immunoblotting a-HuR, followed by goat anti-mouse-HRP (1031-05, Southern Biotech). Cell lysates (1×10^6^ cells/sample) were prepared by standard procedures using RIPA lysis buffer. Proteins were separated on a 4– 12% SDS/PAGE and transferred onto a nitrocellulose membrane by iBlot (Thermo). Mouse monoclonal a-HuR and anti-RhoGDI (MAB9959, Abnova), were used, followed by goat a-rabbit (4050–05) and goat a-mouse-HRP secondary antibodies, respectively (1031-05, both Southern Biotech).

### Flow cytometry and intracellular staining

T cells were washed with FACS buffer (PBS with 1% FBS and 2 mM EDTA) and labeled for 20 min at 4°C with α-CD4 (SK3, BD Horizon), α- CD8 (SK1, BD Horizon). Dead cells were excluded with Near-IR (Life Technologies). For intracellular staining, cells were fixed and permeabilized with Cytofix/Cytoperm kit (BD Biosciences), and stained with α-IFN-γ (4S.B3, BD Bioscience), α-TNF (MAb11, BD Bioscience), α-Puromycin (12D10, Merk). Acquisition was performed using FACS Symphony A5 Cell Analyzer (BD Bioscience). For HuR staining, cells were fixed and permeabilized with eBioscience™ Foxp3/ Transcription Factor Staining Buffer Set (Invitrogen) prior to staining with α-HuR (3A2, Santa Cruz Biotechnology), according to the manufacturer’s protocol. Data were analyzed with FlowJo (BD Biosciences, version 10.8.1).

### Data analysis of flow cytometry data

The FlowJo workspace (.xml or .wsp file) and its FCS files were imported into the R environment (version 4.1.1) (Foundation, R. R: The R Project for Statistical Computing. https://www.r-project.org/.), creating a GatingSet object using CytoML (version 2.4.0) (*67*) . Raw intensity data from gated flow data (IFN-γ, or TNF single positive, and IFN-γ/TNF double positive, or double negative) were extracted from GatingSet using flowWorkspace (version 4.4.0) (*68*) and flowCore (version 2.4.0) (*69*) .

### Data visualization

Results are shown as mean ± SD. Statistical analysis was performed in R- studio, with a two-tailed ratio paired or unpaired Student’s t-test when comparing two groups, or with Krushal-Wallis non-parametric test with Tukey-HSD correction when comparing multiple groups over different time points. p values < 0.05 were considered statistically significant. Data were visualized with ggplot2 (version 3.4.2) (*70*) or using GraphPad Prism (version 9.1.1).

## Acknowledgments

We thank K. Bresser, A.P. Jurgens, and N.D. Zandhuis for critical reading of this manuscript

## Author contributions

Conceptualization, project administration, writing: MVL, ET, MCW. Software, data curation, analysis, visualization: MVL, NS, PS, NK. Validation, investigation, methodology: MVL, BP, AG, AB, LW. Funding acquisition, supervision: MCW.

## Data and materials availability

Code and scripts have been deposited on github (https://github.com/nikolinasostaric/T-cell_smFISH).

## References

1. K. Araki, M. Morita, A. G. Bederman, B. T. Konieczny, H. T. Kissick, N. Sonenberg, R. Ahmed, Translation is actively regulated during the differentiation of CD8 + effector T cells. Nat Immunol 18, 1046–1057 (2017).

2. T. Wolf, W. Jin, G. Zoppi, I. A. Vogel, M. Akhmedov, C. K. E. Bleck, T. Beltraminelli, J. C. Rieckmann, N. J. Ramirez, M. Benevento, S. Notarbartolo, D. Bumann, F. Meissner, B. Grimbacher, M. Mann, A. Lanzavecchia, F. Sallusto, I. Kwee, R. Geiger, Dynamics in protein translation sustaining T cell preparedness. Nat Immunol 21, 927–937 (2020).

3. R. Rak, M. Polonsky, I. Eizenberg-Magar, Y. Mo, Y. Sakaguchi, O. Mizrahi, A. Nachshon, S. Reich-Zeliger, N. Stern-Ginossar, O. Dahan, T. Suzuki, N. Friedman, Y. Pilpel, Dynamic changes in tRNA modifications and abundance during T cell activation. Proc Natl Acad Sci U S A 118, e2106556118 (2021).

4. B. P. Nicolet, M. C. Wolkers, The relationship of mRNA with protein expression in CD8+ T cells associates with gene class and gene characteristics. PLoS One 17 (2022).

5. F. Salerno, N. A. Paolini, R. Stark, M. Von Lindern, M. C. Wolkers, Distinct PKC-mediated posttranscriptional events set cytokine production kinetics in CD8+ T cells. Proc Natl Acad Sci U S A 114, 9677–9682 (2017).

6. B. Popović, B. P. Nicolet, A. Guislain, S. Engels, A. P. Jurgens, N. Paravinja, J. J. Freen-van Heeren, F. P. J. van Alphen, M. van den Biggelaar, F. Salerno, M. C. Wolkers, Time-dependent regulation of cytokine production by RNA binding proteins defines T cell effector function. Cell Rep 42, 112419 (2023).

7. 7. H. Ikeda, L. J. Old, R. D. Schreiber, “The roles of IFN in protection against tumor development and cancer immunoediting” (2002).

8. B. Zhang, T. Karrison, D. A. Rowley, H. Schreiber, IFN-γ- and TNF-dependent bystander eradication of antigen-loss variants in established mouse cancers. Journal of Clinical Investigation 118, 1398–1404 (2008).

9. C. F. Nathan, H. W. Murray, I. E. Wlebe, B. Y. Rubin, Identification of interferon-gamma as the lymphokine that activates human macrophage oxidative metabolism and antimicrobial activity. Journal of Experimental Medicine 158, 670–689 (1983).

10. J. M. Conley, M. P. Gallagher, L. J. Berg, T cells and gene regulation: The switching on and turning up of genes after T cell receptor stimulation in CD8 T cells. Front Immunol 7, 182769 (2016).

11. A. N. Henning, R. Roychoudhuri, N. P. Restifo, Epigenetic control of CD8+ T’cell differentiation. Nature Publishing Group [Preprint] (2018). 10.1038/nri.2017.146.

12. B. Schwanhüusser, D. Busse, N. Li, G. Dittmar, J. Schuchhardt, J. Wolf, W. Chen, M. Selbach, Global quantification of mammalian gene expression control. Nature 2011 473:7347 473, 337–342 (2011).

13. C. Vogel, E. M. Marcotte, Insights into the regulation of protein abundance from proteomic and transcriptomic analyses. Nat Rev Genet 13, 227–232 (2012).

14. F. Salerno, S. Engels, M. van den Biggelaar, F. P. J. van Alphen, A. Guislain, W. Zhao, D. L. Hodge, S. E. Bell, J. P. Medema, M. von Lindern, M. Turner, H. A. Young, M. C. Wolkers, Translational repression of pre-formed cytokine-encoding mRNA prevents chronic activation of memory T cells. Nat Immunol 19, 828– 837 (2018).

15. R. Karki, B. R. Sharma, S. Tuladhar, E. P. Williams, L. Zalduondo, P. Samir, M. Zheng, B. Sundaram, B. Banoth, R. K. S. Malireddi, P. Schreiner, G. Neale, P. Vogel, R. Webby, C. B. Jonsson, T. D. Kanneganti, Synergism of TNF-α and IFN-γ Triggers Inflammatory Cell Death, Tissue Damage, and Mortality in SARS-CoV-2 Infection and Cytokine Shock Syndromes. Cell 184, 149–168.e17 (2021).

16. K. P. Hoefig, A. Reim, C. Gallus, E. H. Wong, G. Behrens, C. Conrad, M. Xu, L. Kifinger, T. Ito-Kureha, K. A. Y. Defourny, A. Geerlof, J. Mautner, S. M. Hauck, D. Baumjohann, R. Feederle, M. Mann, M. Wierer, E. Glasmacher, V. Heissmeyer, Defining the RBPome of primary T helper cells to elucidate higher-order Roquin-mediated mRNA regulation. Nature Communications 2021 12:1 12, 1–18 (2021).

17. J. I. Perez-Perri, B. Rogell, T. Schwarzl, F. Stein, Y. Zhou, M. Rettel, A. Brosig, M. W. Hentze, Discovery of RNA-binding proteins and characterization of their dynamic responses by enhanced RNA interactome capture. Nature Communications 2018 9:1 9, 1–13 (2018).

18. K. Deka, S. Saha, Heat stress induced arginylation of HuR promotes alternative polyadenylation of Hsp70.3 by regulating HuR stability and RNA binding. Cell Death & Differentiation 2020 28:2 28, 730–747 (2020).

19. M. D. Diaz-Muñoz, S. E. Bell, K. Fairfax, E. Monzon-Casanova, A. F. Cunningham, M. Gonzalez-Porta, S. R. Andrews, V. I. Bunik, K. Zarnack, T. Curk, W. A. Heggermont, S. Heymans, G. E. Gibson, D. L. Kontoyiannis, J. Ule, M. Turner, The RNA-binding protein HuR is essential for the B cell antibody response. Nature Immunology 2015 16:4 16, 415–425 (2015).

20. I. C. Osma-Garcia, D. Capitan-Sobrino, M. Mouysset, S. E. Bell, M. Lebeurrier, M. Turner, M. D. Diaz-Muñoz, The RNA-binding protein HuR is required for maintenance of the germinal centre response. Nat Commun 12 (2021).

21. J. R. Poganik, M. J. C. Long, M. T. Disare, X. Liu, S. H. Chang, T. Hla, Y. Aye, Post-transcriptional regulation of Nrf2-mRNA by the mRNA-binding proteins HuR and AUF1. FASEB Journal 33, 14636–14652 (2019).

22. C. Tiedje, N. Ronkina, M. Tehrani, S. Dhamija, K. Laass, H. Holtmann, A. Kotlyarov, M. Gaestel, The p38/MK2-Driven Exchange between Tristetraprolin and HuR Regulates AU-Rich Element-Dependent Translation. PLoS Genet 8 (2012).

23. F. Salerno, M. Turner, M. C. Wolkers, Dynamic Post-Transcriptional Events Governing CD8+ T Cell Homeostasis and Effector Function. Elsevier Ltd [Preprint] (2020). 10.1016/j.it.2020.01.001.

24. N. D. Zandhuis, B. P. Nicolet, M. C. Wolkers, RNA-Binding Protein Expression Alters Upon Differentiation of Human B Cells and T Cells. Front Immunol 12, 717324 (2021).

25. S. Das, M. Vera, V. Gandin, R. H. Singer, E. Tutucci, Intracellular mRNA transport and localized translation. Nat Rev Mol Cell Biol 22, 483–504 (2021).

26. K. Davari, J. Lichti, C. Gallus, F. Greulich, N. H. Uhlenhaut, M. Heinig, C. C. Friedel, E. Glasmacher, Rapid Genome-wide Recruitment of RNA Polymerase II Drives Transcription, Splicing, and Translation Events during T Cell Responses. Cell Rep 19, 643–654 (2017).

27. F. Radford, S. Tyagi, M. L. Gennaro, R. Pine, Y. Bushkin, Flow Cytometric Characterization of Antigen- Specific T Cells Based on RNA and Its Advantages in Detecting Infections and Immunological Disorders. Critical Reviews&trade; in Immunology 36, 359–378 (2016).

28. B. P. Nicolet, A. Guislain, M. C. Wolkers, Combined Single-Cell Measurement of Cytokine mRNA and Protein Identifies T Cells with Persistent Effector Function. The Journal of Immunology 198, 962–970 (2017).

29. Q. Han, N. Bagheri, E. M. Bradshaw, D. A. Hafler, D. A. Lauffenburger, J. C. Love, Polyfunctional responses by human T cells result from sequential release of cytokines. Proc Natl Acad Sci U S A 109, 1607– 1612 (2012).

30. A. M. Femino, F. S. Fay, K. Fogarty, R. H. Singer, Visualization of single RNA transcripts in situ. Science 280, 585–590 (1998).

31. A. Raj, P. van den Bogaard, S. A. Rifkin, A. van Oudenaarden, S. Tyagi, Imaging individual mRNA molecules using multiple singly labeled probes. Nat Methods 5, 877–879 (2008).

32. M. Fang, H. Xie, S. K. Dougan, H. Ploegh, A. van Oudenaarden, Stochastic Cytokine Expression Induces Mixed T Helper Cell States. PLoS Biol 11, e1001618 (2013).

33. A. Senecal, B. Munsky, F. Proux, N. Ly, F. E. Braye, C. Zimmer, F. Mueller, X. Darzacq, Transcription Factors Modulate c-Fos Transcriptional Bursts. Cell Rep 8, 75–83 (2014).

34. Y. Bushkin, F. Radford, R. Pine, A. Lardizabal, B. T. Mangura, M. L. Gennaro, S. Tyagi, Profiling T cell activation using single molecule-FISH and flow cytometry. J Immunol 194, 836 (2014).

35. D. M. Franchini, O. Lanvin, M. Tosolini, E. Patras de Campaigno, A. Cammas, S. Péricart, C. M. Scarlata, M. Lebras, C. Rossi, L. Ligat, F. Pont, P. B. Arimondo, C. Laurent, M. Ayyoub, F. Despas, M. Lapeyre- Mestre, S. Millevoi, J. J. Fournié, Microtubule-Driven Stress Granule Dynamics Regulate Inhibitory Immune Checkpoint Expression in T Cells. Cell Rep 26, 94–107.e7 (2019).

36. W. Ma, C. Mayr, A Membraneless Organelle Associated with the Endoplasmic Reticulum Enables 3’UTR- Mediated Protein-Protein Interactions. Cell 175, 1492–1506.e19 (2018).

37. R. Chouaib, A. Safieddine, X. Pichon, A. Imbert, O. S. Kwon, A. Samacoits, A. M. Traboulsi, M. C. Robert, N. Tsanov, E. Coleno, I. Poser, C. Zimmer, A. Hyman, H. Le Hir, K. Zibara, M. Peter, F. Mueller, T. Walter, E. Bertrand, A Dual Protein-mRNA Localization Screen Reveals Compartmentalized Translation and Widespread Co-translational RNA Targeting. Dev Cell 54, 773–791.e5 (2020).

38. E. Tutucci, A. Maekiniemi, J. L. Snoep, M. Seiler, K. Van Rossum, D. D. Van Niekerk, P. Savakis, K. Zarnack, R. H. Singer, Cyclin CLB2 mRNA localization determines efficient protein synthesis 1 to orchestrate bud growth and cell cycle progression 2. doi: 10.1101/2022.03.01.481833.

39. Y. Bushkin, F. Radford, R. Pine, A. Lardizabal, B. T. Mangura, M. L. Gennaro, S. Tyagi, Profiling T Cell Activation Using Single-Molecule Fluorescence In Situ Hybridization and Flow Cytometry. The Journal of Immunology 194, 836–841 (2015).

40. C. Eliscovich, S. M. Shenoy, R. H. Singer, Imaging mRNA and protein interactions within neurons. Proc Natl Acad Sci U S A 114, E1875–E1884 (2017).

41. F. Mueller, A. Senecal, K. Tantale, H. Marie-Nelly, N. Ly, O. Collin, E. Basyuk, E. Bertrand, X. Darzacq, C. Zimmer, FISH-quant: Automatic counting of transcripts in 3D FISH images. [Preprint] (2013). 10.1038/nmeth.2406.

42. A. Maekiniemi, R. H. Singer, E. Tutucci, Single molecule mRNA fluorescent in situ hybridization combined with immunofluorescence in S. cerevisiae: Dataset and quantification. Data Brief 30, 105511 (2020).

43. S. van Otterdijk, M. Motealleh, Z. Wang, T. D. Visser, P. Savakis, E. Tutucci, Single-Molecule Fluorescent In Situ Hybridization (smFISH) for RNA Detection in the Fungal Pathogen Candida albicans. Methods Mol Biol 2784, 25–44 (2024).

44. C. Stringer, T. Wang, M. Michaelos, M. Pachitariu, Cellpose: a generalist algorithm for cellular segmentation. Nature Methods 2020 18:1 18, 100–106 (2020).

45. J. R. Almeida, D. A. Price, L. Papagno, Z. A. Arkoub, D. Sauce, E. Bornstein, T. E. Asher, A. Samri, A. Schnuriger, I. Theodorou, D. Costagliola, C. Rouzioux, H. Agut, A. G. Marcelin, D. Douek, B. Autran, V. Appay, Superior control of HIV-1 replication by CD8+ T cells is reflected by their avidity, polyfunctionality, and clonal turnover. J Exp Med 204, 2473–2485 (2007).

46. 46. R. De Groot, M. M. Van Loenen, A. Guislain, B. P. Nicolet, J. J. Freen-Van Heeren, O. J. H. M. Verhagen, M. M. Van Den Heuvel, J. De Jong, P. Burger, C. E. Van Der Schoot, R. M. Spaapen, D. Amsen, J. B. A. G. Haanen, K. Monkhorst, K. J. Hartemink, M. C. Wolkers, Polyfunctional tumor-reactive T cells are effectively expanded from non-small cell lung cancers, and correlate with an immune-engaged T cell profile. Oncoimmunology 8 (2019).

47. I. Horvathova, F. Voigt, A. V. Kotrys, Y. Zhan, C. G. Artus-Revel, J. Eglinger, M. B. Stadler, L. Giorgetti, J. A. Chao, The Dynamics of mRNA Turnover Revealed by Single-Molecule Imaging in Single Cells. Mol Cell 68, 615–625.e9 (2017).

48. A. C. Tuck, A. Rankova, A. B. Arpat, L. A. Liechti, D. Hess, V. Iesmantavicius, V. Castelo-Szekely, D. Gatfield, M. Bühler, Mammalian RNA Decay Pathways Are Highly Specialized and Widely Linked to Translation. Mol Cell 77, 1222–1236.e13 (2020).

49. K. Rothamel, S. Arcos, B. Kim, C. Reasoner, S. Lisy, N. Mukherjee, M. Ascano, ELAVL1 primarily couples mRNA stability with the 3′ UTRs of interferon-stimulated genes. Cell Rep 35, 109178 (2021).

50. W. Wang, M. C. Caldwell, S. Lin, H. Furneaux, M. Gorospe, HuR regulates cyclin A and cyclin B1 mRNA stability during cell proliferation. EMBO J 19, 2340–2350 (2000).

51. Y. Akaike, K. Masuda, Y. Kuwano, K. Nishida, K. Kajita, K. Kurokawa, Y. Satake, K. Shoda, I. Imoto, K. Rokutan, HuR Regulates Alternative Splicing of the TRA2β Gene in Human Colon Cancer Cells under Oxidative Stress. Mol Cell Biol 34, 2857–2873 (2014).

52. L. S. Namer, F. Osman, Y. Banai, B. Masquida, R. Jung, R. Kaempfer, An Ancient Pseudoknot in TNF-α Pre-mRNA Activates PKR, Inducing eIF2α Phosphorylation that Potently Enhances Splicing. Cell Rep 20, 188–200 (2017).

53. 53. B. Yang Yang, J.-F. Chang, J. R. Parnes, C. Garrison Fathman, “T Cell Receptor (TCR) Engagement Leads to Activation-induced Splicing of Tumor Necrosis Factor (TNF) Nuclear Pre-mRNA” (1998); http://www.jem.org.

54. H. A. Meijer, M. Bushell, K. Hill, T. W. Gant, A. E. Willis, P. Jones, C. H. de Moor, A novel method for poly(A) fractionation reveals a large population of mRNAs with a short poly(A) tail in mammalian cells. Nucleic Acids Res 35, e132 (2007).

55. N. Battich, T. Stoeger, L. Pelkmans, Control of Transcript Variability in Single Mammalian Cells. Cell 163, 1596–1610 (2015).

56. B. C. Mercier, E. Labaronne, D. Cluet, L. Guiguettaz, N. Fontrodona, A. Bicknell, A. Corbin, M. Wencker, F. Aube, L. Modolo, K. Jouravleva, D. Auboeuf, M. J. Moore, E. P. Ricci, Translation-dependent and - independent mRNA decay occur through mutually exclusive pathways defined by ribosome density during T cell activation. Genome Res 34, 394–409 (2024).

57. L. A. Passmore, J. Coller, Roles of mRNA poly(A) tails in regulation of eukaryotic gene expression. Nature Reviews Molecular Cell Biology 2021 23:2 23, 93–106 (2021).

58. W. Dai, G. Zhang, E. V. Makeyev, RNA-binding protein HuR autoregulates its expression by promoting alternative polyadenylation site usage. Nucleic Acids Res 40, 787–800 (2012).

59. J. Cherry, H. Jones, V. A. Karschner, P. H. Pekala, Post-transcriptional Control of CCAAT/Enhancer-binding Protein β (C/EBPβ) Expression: FORMATION OF A NUCLEAR HuR-C/EBPβ mRNA COMPLEX DETERMINES THE AMOUNT OF MESSAGE REACHING THE CYTOSOL. Journal of Biological Chemistry 283, 30812–30820 (2008).

60. W. Zhang, A. C. Vreeland, N. Noy, RNA-binding protein HuR regulates nuclear import of protein. J Cell Sci 129, 4025 (2016).

61. E. L. Van Nostrand, P. Freese, G. A. Pratt, X. Wang, X. Wei, R. Xiao, S. M. Blue, J. Y. Chen, N. A. L. Cody, D. Dominguez, S. Olson, B. Sundararaman, L. Zhan, C. Bazile, L. P. B. Bouvrette, J. Bergalet, M. O. Duff, K. E. Garcia, C. Gelboin-Burkhart, M. Hochman, N. J. Lambert, H. Li, M. P. McGurk, T. B. Nguyen, T. Palden, I. Rabano, S. Sathe, R. Stanton, A. Su, R. Wang, B. A. Yee, B. Zhou, A. L. Louie, S. Aigner, X. D. Fu, E. Lécuyer, C. B. Burge, B. R. Graveley, G. W. Yeo, A large-scale binding and functional map of human RNA-binding proteins. Nature 2020 583:7818 583, 711–719 (2020).

62. E. J. Wherry, S. J. Ha, S. M. Kaech, W. N. Haining, S. Sarkar, V. Kalia, S. Subramaniam, J. N. Blattman, D. L. Barber, R. Ahmed, Molecular Signature of CD8+ T Cell Exhaustion during Chronic Viral Infection. Immunity 27, 670–684 (2007).

63. X. Hu, L. B. Ivashkiv, Cross-regulation of Signaling Pathways by Interferon-γ: Implications for Immune Responses and Autoimmune Diseases. [Preprint] (2009). 10.1016/j.immuni.2009.09.002.

64. D. I. Jang, A. H. Lee, H. Y. Shin, H. R. Song, J. H. Park, T. B. Kang, S. R. Lee, S. H. Yang, The role of tumor necrosis factor alpha (Tnf-α) in autoimmune disease and current tnf-α inhibitors in therapeutics. MDPI AG [Preprint] (2021). 10.3390/ijms22052719.

65. M. Vera, E. Tutucci, R. H. Singer, Imaging Single mRNA Molecules in Mammalian Cells Using an Optimized MS2-MCP System. Methods in Molecular Biology 2038, 3–20 (2019).

66. S. Rozen, H. Skaletsky, Primer3 on the WWW for General Users and for Biologist Programmers. Methods Mol Biol 132, 365–386 (2000).

67. G. Finak, W. Jiang, R. Gottardo, CytoML for cross-platform cytometry data sharing. Cytometry Part A 93, 1189–1196 (2018).

68. 68. A. Greg Finak, M. Jiang Maintainer Greg Finak, M. Jiang, Title Infrastructure for representing and interacting with gated and ungated cytometry data sets. (2024).

69. F. Hahne, N. LeMeur, R. R. Brinkman, B. Ellis, P. Haaland, D. Sarkar, J. Spidlen, E. Strain, R. Gentleman, flowCore: A Bioconductor package for high throughput flow cytometry. BMC Bioinformatics 10, 1–8 (2009).

70. H. Wickham, ggplot2. doi: 10.1007/978-3-319-24277-4 (2016).

