## Supplementary Material and Methods, Supplementary figures and Tables for "Single molecule imaging of transcription dynamics, RNA localization and fate in human T cells"

Maria Valeria Lattanzio *et al.*

**This PDF file includes:**

Supplementary Materials and Methods

Figs. S1 to S5

Tables S1-S3

Movies S1 to S2

**Other Supplementary Materials for this manuscript include the following:**

Movies S1 to S2

**Suppl. Material and Methods**

**Coverslip coating selection**

For efficient T cell attachment, we compared different coating methods (**Suppl. MeM Fig.A)**: CD8^+^ T cells were coated on round 16-mm coverslips (Fisherbrand Borosilicate Glass Circle Coverslip) using (**2**) Cytospin (3min x1800rpm), or on coverslips coated with (**2**) Poly-L-lysine high molecular weight (0.1 % (w/v) in H_2_O, Sigma), **(3)** Poly-L-lysine low molecular weight (0.1 % (w/v) in H_2_O, Sigma) (**4**), Corning^®^ Cell-Tak(TM) Cell (**1**) and (**5**) Tissue Adhesive and Alcian Blue (Alcian blue in 3% acetic acid, Sigma). Corning® Cell-Tak™ (**1**) and mechanical seeding using cytospin (**2**) resulted in high-density seeding, but perturbed the shape of T cells. Coating with Poly-L-lysine (high MW (**3**) and low MW (**4**)) resulted in poor T cell attachment. With high T cell attachment and maintaining T cell shape, Alcian Blue (**5**) was most optimal for mounting T cells.

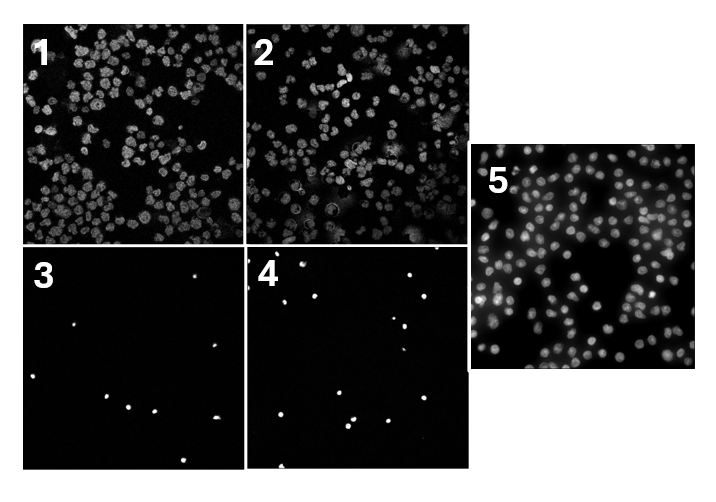
**Suppl. MeM Fig.A**

**Suppl. MeM Fig.A** Optimizing Coverslip coating. MAX projection of resting CD3^+^ T cells on coverslips mechanically seeded with cytospin (2), or on cover slips coated with Corning® Cell-Tak™ Cell and Tissue Adhesive (1), Poly-L-lysine high MW (3), Poly-L-lysine MW (4), or Alcian Blue (5). T cells were stained with DAPI.

**T cell amount for coating**

To obtain sufficient CD8^+^ T cells for smFISH measurements, we tested different numbers of CD8^+^ T cells mounted onto Alcian Blue coated coverslips (**Suppl.MeM Fig.B**).This titration essay showed that 1x10^6^ T cells was most optimal.

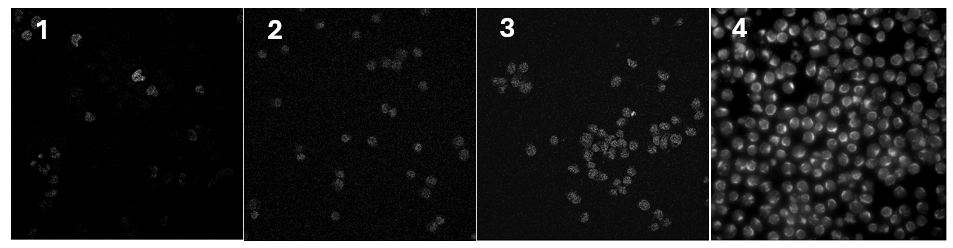
**Suppl.MeM Fig.B**

**Suppl.MeM Fig.B Optimizing cell numbers for coating.** MAX projection of resting CD3+ T cells on coverslips coated with Alcian Blue using different amount of cells: 250.000 cells (1), 500.000 cells (2), 750.000 cells (3) and 1x10^6^ T cells (4). The images are MAX projection of cellular autofluorescence in FITC channel

**Fluorophore and filter setting selection**

To obtain a high-quality smFISH signal, low background signal from cells is key. We found that autofluorescence signal of CD3^+^ T cells presented with a dot shape (**Suppl.MeM Fig.C;** see white arrows**)** in the FITC channel (**1**), Cy3 channel (**2**) and Cy5 channel (**3**). While these autofluorescence dots do not interfere with standard (protein) immunofluorescence, they can be mislabelled as single-molecule RNA. We therefore selected Cal-Fluo 610 (Cy3.5 channel) and Quasar-670 (Cy5 channel) as fluorophores conjugated to *IFNG* and *TNF* smFISH probes, respectively, and we used the AHF-LED-FISH-R Filter for Cy3.5 and the F36-523 Cy5 HCBrightLine Filter for Cy5 to reduce the background signal to increase the signal-to-noise ratio.

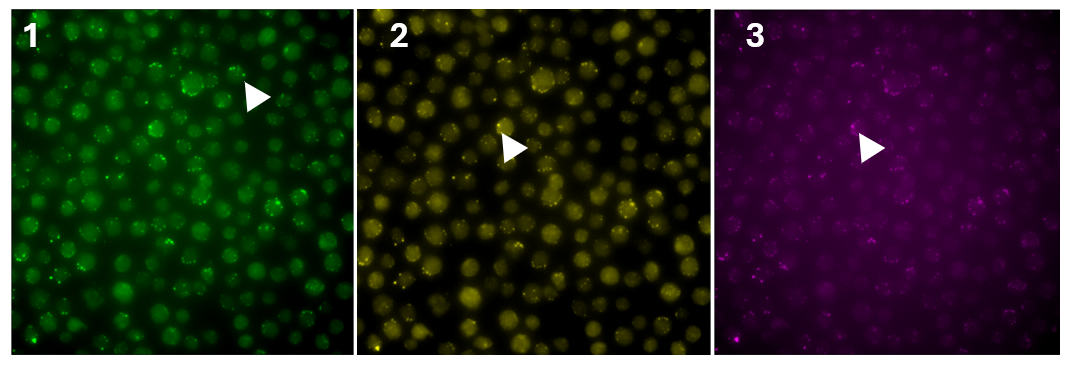
**Suppl.MeM Fig.C**

**Suppl. MeM Fig.C Autofluorescence of unstained resting CD3^+^ T cells.** MAX projection of FITC channel (**1**), Cy3 (**2**) and Cy5 (**3**). White arrows indicate the autofluorescence signal in structures of dots.

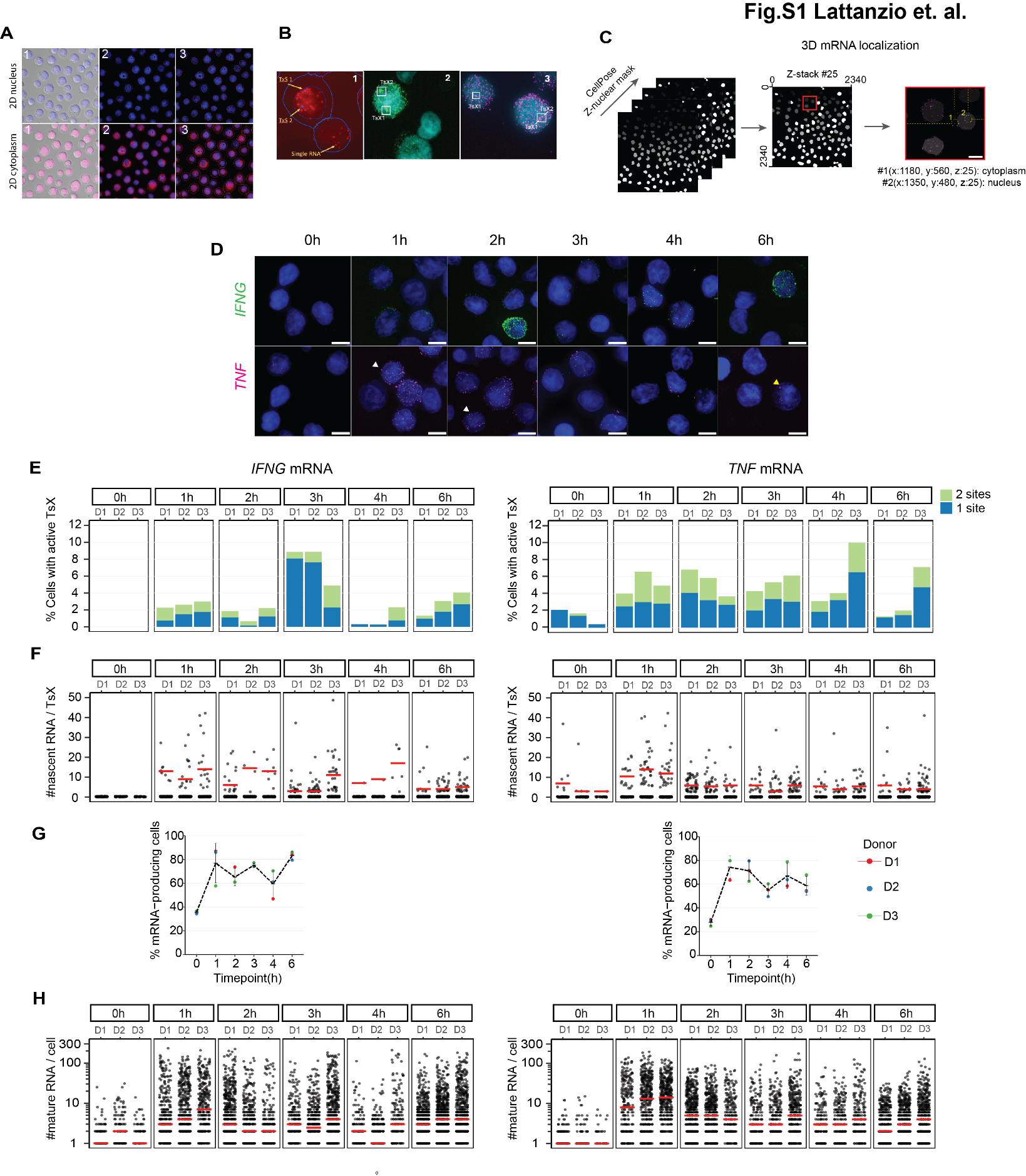
Fig. S1.

Fig. S1. *IFNG* and *TNF* mRNA quantification with T-cell smFISH

**A.** Schematic representation of 2D workflow to define nucleus (top panel) and cytoplasm (bottom panel) with fq-segmentation (FISH-quant). (1) Merged MAX projection of DIC with DAPI (nucleus, blue) or autofluorescence in the Quasar-670 channel (cell mask, magenta). (2) MAX projection of DAPI (top) and Quasar-670 (bottom) as input for fq-segmentation. (3) Nuclear (top) and cellular (bottom) outline depicted from fq-segmentation, indicated with red line. **B.** Maximal projection of CD8^+^ T cells activated for 1h with α-CD3/α-CD28. Example of TsX identification and single mRNA using FISH-Quant Matlab (**1**) (blue line: 2D cellular outline, dotted blue line: 2D nuclear outline). Example of *IFNG* mRNA (green, **2**) and *TNF* mRNA (magenta,**3**) smFISH merged on DAPI staining (cyan). Boxes indicate the number of Transcription site (TsX1 or TsX 2). **C.** T-cell smFISH analysis pipeline. Z-stack nuclear masks (indicated in shades of gray) obtained with CellPose nuclear mask function. Pixel coordinates are extracted (range 0, 2340 pixels) per Z-nuclear mask. Mask #25 as an example. T-cell smFISH analysis extracts coordinate information to define localization of mRNA molecules inside or outside the nuclear/cytoplasmic mask, as exemplified for mRNA molecules in yellow dotted circles: mRNA#1: cytoplasmic, mRNA#2: nuclear. Scale bar: 5μm. **D.** Maximal projection of DAPI (blue), *IFNG* mRNA (top panel), and *TNF* mRNA (bottom panel) in Teff cells re-activated with α-CD3/α-CD28. White arrows: single mRNA molecules. Yellow arrows: TsX. Scale bar: 5 μm. **E.** Percentage of Teff cells with 1 (blue) or 2 (green) active transcription sites (TsX) for *IFNG* (left) and *TNF* (right) after α-CD3/α-CD28 stimulation. N=3 donors (D). **F.** Number of nascent *IFNG* (left) and *TNF* (right) RNA per active TsX. Each dot represents one TsX. n=3 donors. Red bar: median. **G.** Percentage of Teff cells expressing mature mRNA. n=3 donors, line: mean. **H.** Number of mature *IFNG* (left panel) and *TNF* (right panel) mRNA of Teff expressing ≥ 1 mature mRNA. Each dot represents one cell. n=3 donors. Red bar: median.

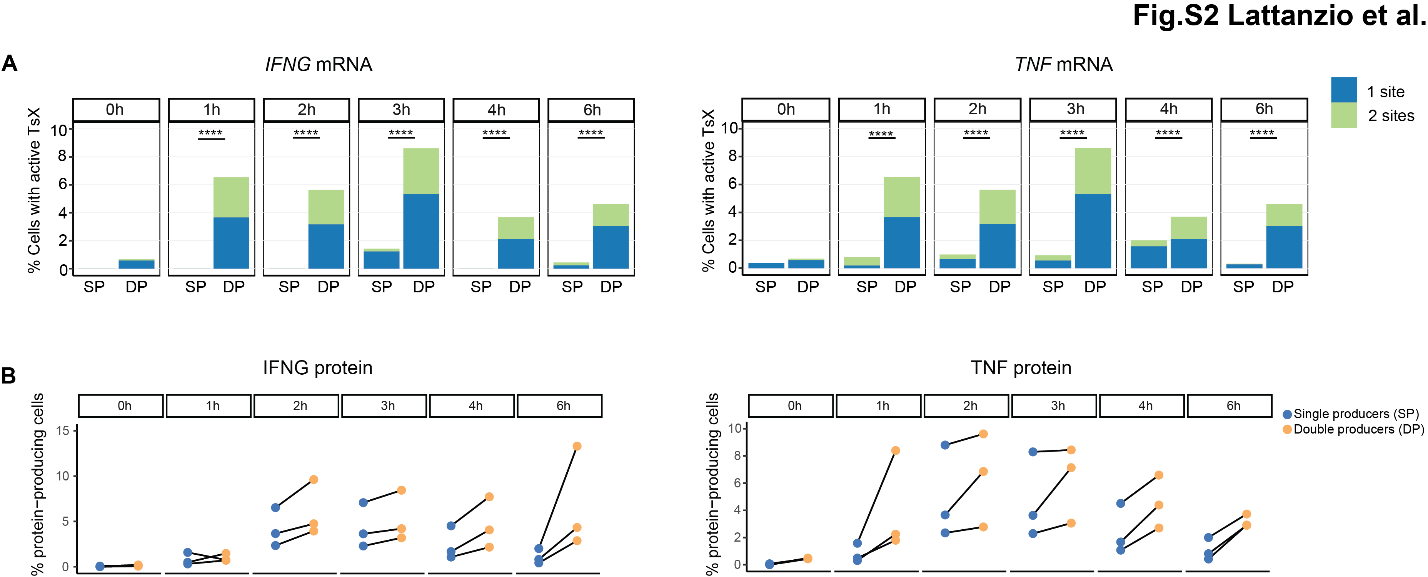
Fig. S2.

Fig. S2. *IFNG* and *TNF* transcription in single and double positive expressors

**A.** Percentage of Teff cells with one (SP, blue) or two (DP, green) active transcription sites for *IFNG* (left) and *TNF* (right) upon α-CD3/α-CD28 stimulation. Data depict median of 3 pooled donors. *p≤0.05, **p≤0.01, ***p≤0.001, ****p≤0.0001 ns: non-significant. Kruskal-Wallis non-parametric test, and post-hoc Tukey HSD test. **B.** IFN-γ (left) and TNF (right) protein expression of SP (blue) and DP (orange) protein-producing Teff cells. Bref A was added for a maximum of 2h of activation. Each dot indicates 1 donor.

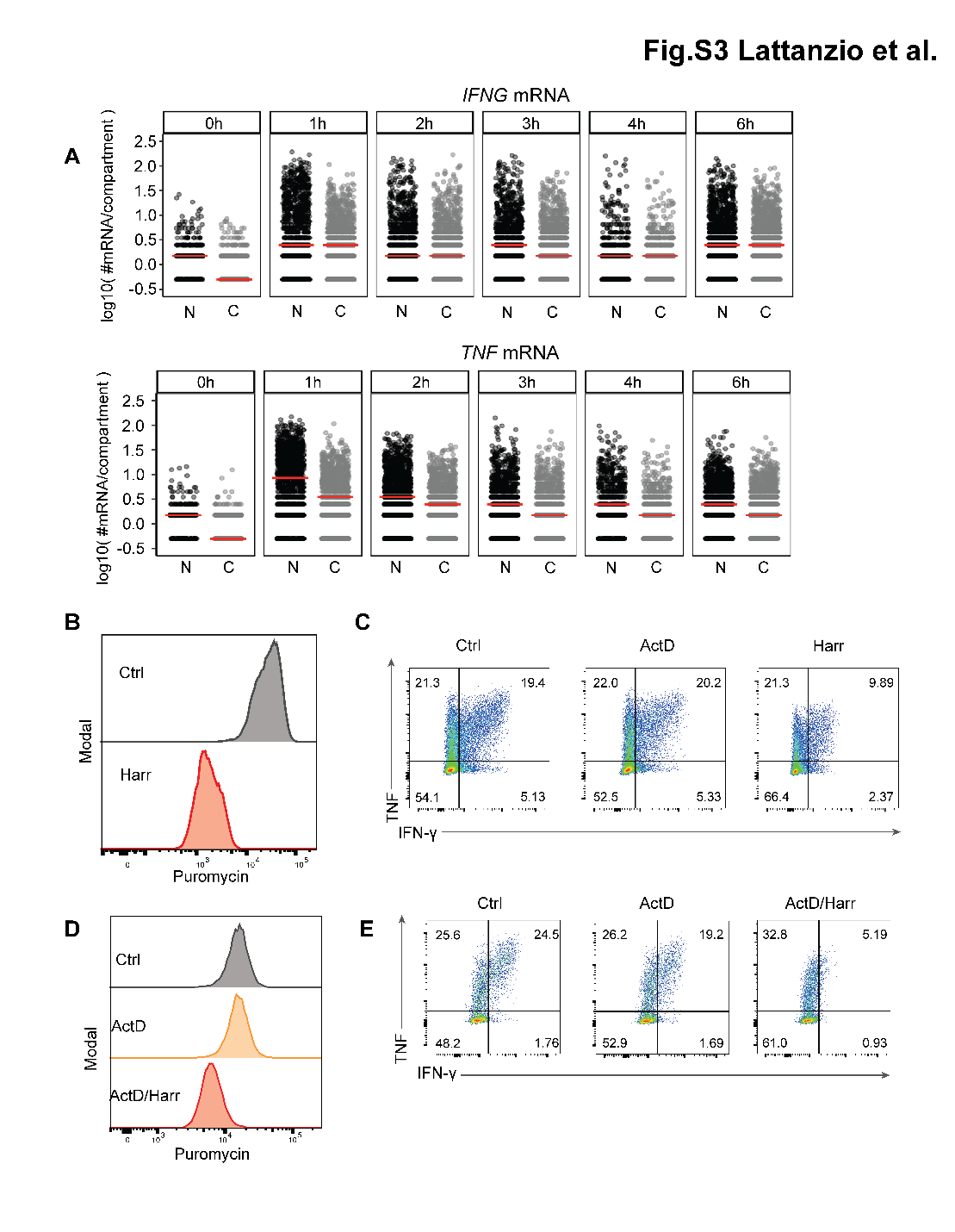
Fig. S3.

**Fig. S3. IFNG and TNF expression depends on translation control**

**A.** Number of nuclear (N, grey) and cytoplasmic (C, black) cytokine mRNA in Teff cells expressing ≥ 1 mature RNA. Data representation in log10 scale (y-axis), with a coefficient of 0.5 added. Red bar: median **B.** Puromycin expression in Teff cells activated for 2h with α-CD3/α-CD28 that were treated for the last hour with Harringtonine (HARR) or left untreated (Ctrl). Puromycin was added for the last 10 min of T cell activation **B.** IFN-γ and TNF protein expression in Teff cells activated for 2h with α-CD3/α-CD28 that were treated for the 2^nd^ hour with Actinomycin D (ActD), HARR, or left untreated. **C.** Puromycin expression in Teff cells activated for 2h with α-CD3/α-CD28 that were treated in the 2^nd^ hour with indicated drug or left untreated (Ctrl). Depicted data representative for 3 donors. **D.** IFN-γ and TNF protein expression of Teff cells activated for 2h and treated in the 2^nd^ hour with indicated drug. Data representative of 3 donors.

Fig. S4.

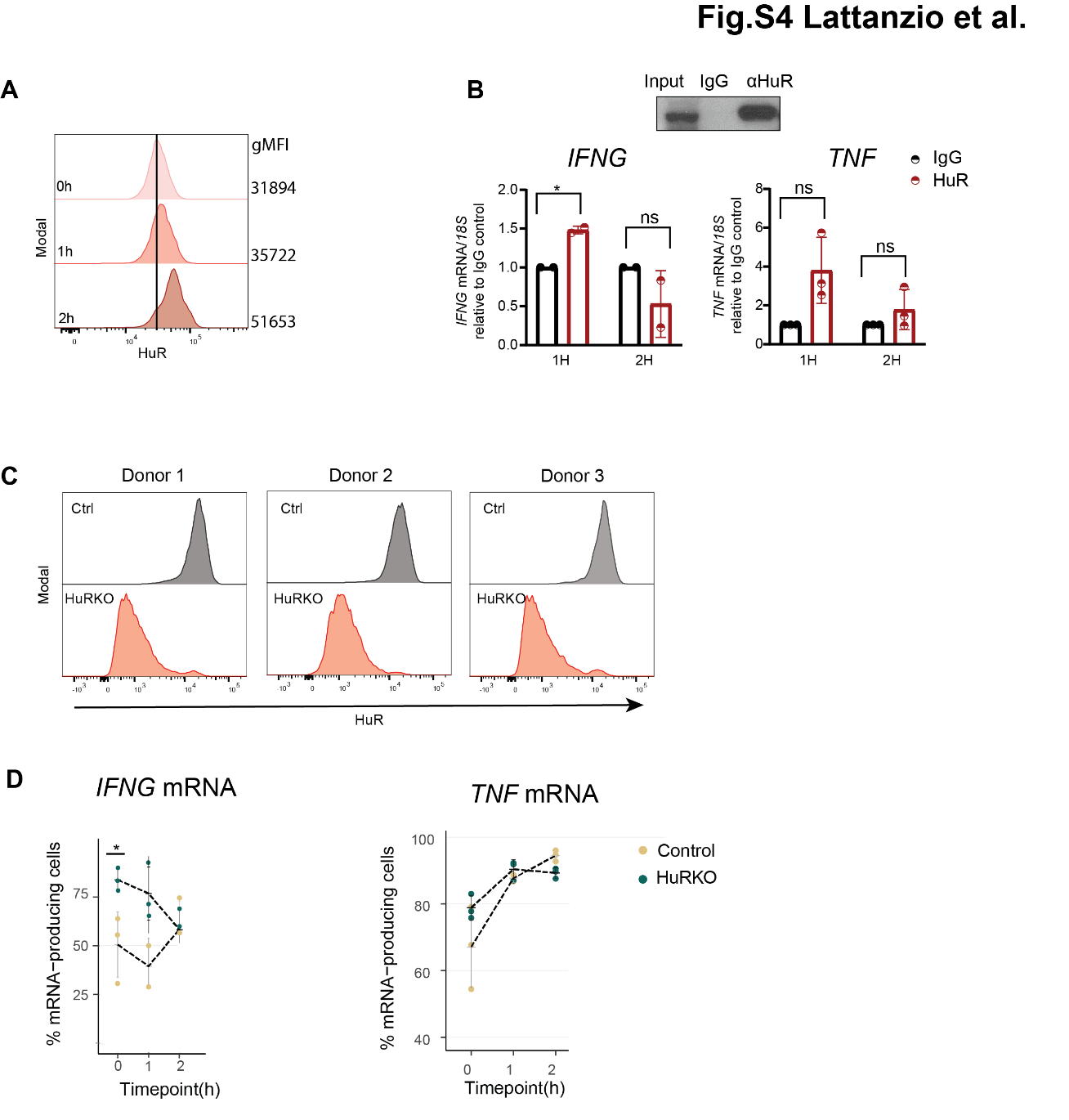

**Fig. S4.** **Expression and interaction kinetics of HuR with *IFNG* and *TNF* mRNA**

**A.** HuR protein expression in Teff cells that were resting (0h), or that were activated with α-CD3/α-CD28 for indicated time points. Geometric mean fluorescence intensity (gMFI) of HuR protein expression indicated on the right. Data representative of 3 donors. **B.** Native RNA immunoprecipitation (RIP) with α-HuR or immunoglobulin G (IgG) isotype control from Teff cells activated for 1h and 2h with PMA-ionomycin. (Top) Immunoblot of HuR expresssion upon RIP. (Bottom) qRT-PCR of endogenous *IFNG* or *TNF* mRNA from RIP with HuR or IgG control antibodies. Data compiled of two (*IFNG*) or three (*TNF*) donors from independently performed experiments. Mean ± SD. *p≤0.05, **p≤0.01, ***p≤0.001, ****p≤0.0001 ns: non-significant. Two-tailed paired t-test. **C.** HuR expression in HuR-KO and control-nucleofected (Ctrl) Teff cells by flow cytometry. n=3 donors. **D.** Percentage of Teff cells expressing ≥1 mature *IFNG* (left) and *TNF* (right) mRNA in Control and HuR KO Teff cells. Each dot indicates 1 donor. Line: mean. (*p≤0.05, post-hoc Tukey HSD).

Fig. S5.

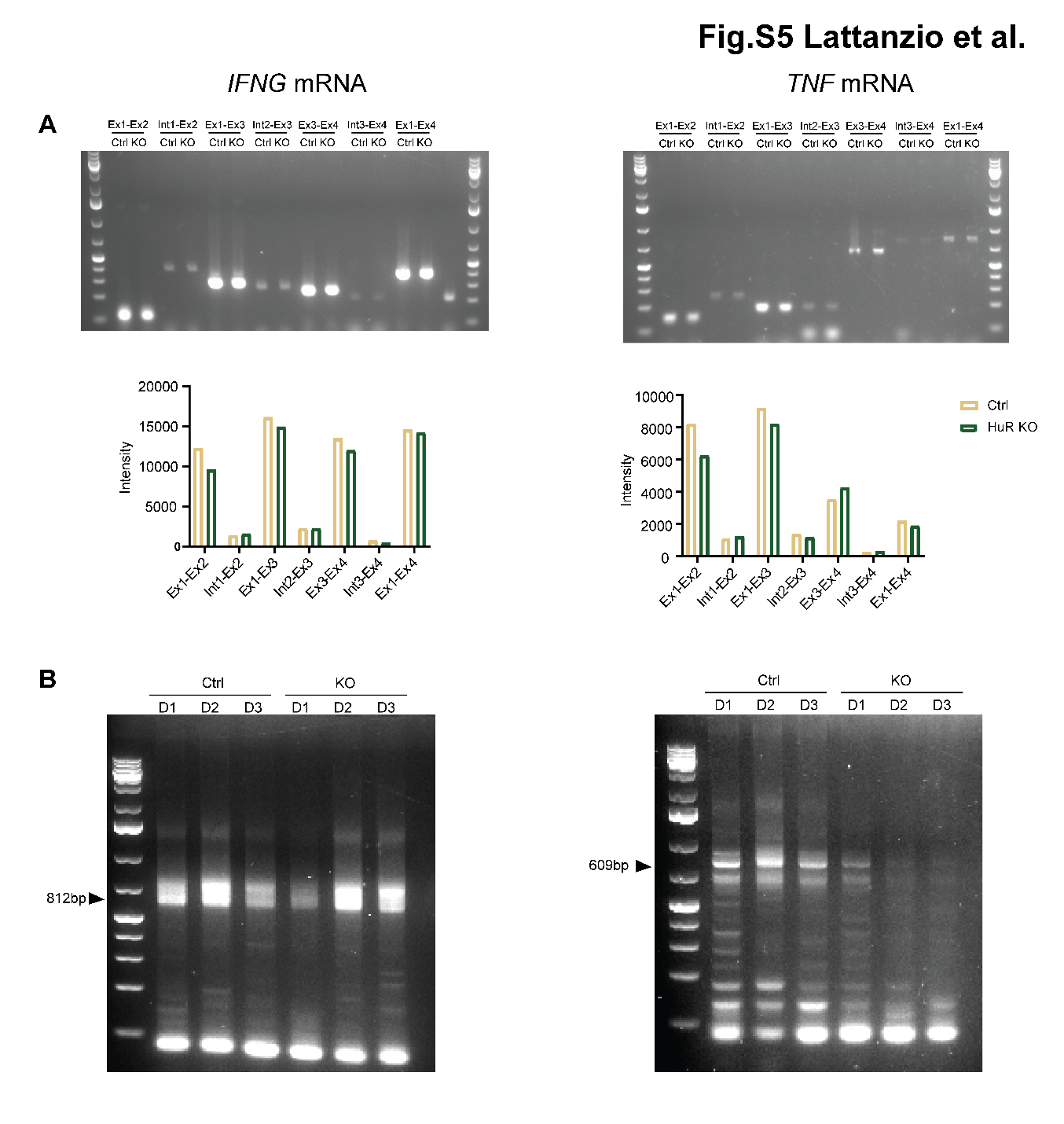
**Fig. S5.** **Analysis of *IFNG* and *TNF* splicing and polyadenylation**

**A.** Measurement of *IFNG* (left) and *TNF* (right) intron-exon junctions with RT-PCR of Control (Ctrl) and HuR KO (KO) Teff cells that were stimulated for 1h with α-CD3/α-CD28. Top: RT-PCR gel. Bottom: quantification of band intensity. **B.** Measurement of poly(A) length of cytokine mRNA in control (Ctrl) or HuR-KO (KO) Teff cells activated for 1h with α-CD3/α-CD28 using RL-PAT assay. n=3 donors. Arrows indicate *IFNG* 3’UTR full-length: 812bp, and *TNF* 3’UTR full-length: 609bp.

Table S1.

***IFNG* and *TNF* probes for T cell smFISH**

| **human *IFNG* probes** | **human *TNF* probes** |
| --- | --- |
| agctgatcttcagatgatca | ggtctgtagttgcttctctc |
| caggtccaaaggacttaact | cgtctgagggttgttttcag |
| gtagttcttgtatcaagctg | gggtcagtatgtgagaggaa |
| atttcatcgtttccgagaga | atcatgctttcagtgctcat |
| gcagagctgaaaagccaaga | aggctgaggaacaagcaccg |
| taacagccaagagaacccaa | tgccacgatcaggaaggaga |
| ttttacatatgggtcctggc | caaagtgcagcaggcagaag |
| atttcttaaggttttctgct | gagggctgattagagagagg |
| catctgaatgacctgcatta | cggggttcgagaagatgatc |
| aaaagagttccattatccgc | cttgagggtttgctacaaca |
| tatttttctgtcactctcct | accagctggttatctctcag |
| aggagacaatttggctctgc | ctgggagtagatgaggtaca |
| gctctggtcatctttaaagt | ctgatggtgtgggtgaggag |
| atggtctccacactcttttg | tctggtaggagacggcgatg |
| acttgacattcatgtcttcc | ctcttgatggcagagaggag |
| tcgtttctttttgttgctat | gatagatgggctcataccag |
| tagtcagcttttcgaagtca | attgatctcagcgctgagtc |
| attcaagtcagttaccgaat | caaagtcgagatagtcgggc |
| catgtattgctttgcgttgg | aaagtagacctgcccagact |
| tcagccatcacttggatgag | ctcctcacagggcaatgatc |
| tgttttagctgctggcgaca | ttgggaaggttggatgttcg |
| atctgactcctttttcgctt | ataaagggattggggcaggg |
| atgctcttcgacctcgaaac | ttgagggtgtctgaaggagg |
| aggcaggacaaccattactg | tctctttttgagccagaaga |
| agtgagacagtcacaggata | aagttctaagcttgggttcc |
| acatagccttgcctaattag | tcgaagtggtggtcttgttg |
| ccctgagataaagccttgta | cacacattcctgaatcccag |
| ttaggttggctgcctagttg | ttgaattcttagtggttgcc |
| aaacacacaacccatgggat | agggatcaaagctgtaggcc |
| gttcattgtatcatcaagtg | tggtctccagattccagatg |
| ctggatagtatcacttcact | cattctggccagaaccaaag |
| gcatattttcaaaccggcag | taggtgaggtcttctcaagt |
| aagttctgtctgacatgcca | aaggtccacttgtgtcaatt |
| atcagggtcacctgacacat | acatctggagagaggaaggc |
| tctcctgagatgctatgttt | cgtgtctcaaggaagtctgg |
| tttggaagcaccaggcatga | taaatagagggagctggctc |
| atgagttactttccatttgg | ccggtctcccaaataaatac |
|  | caaggcagctcctacattgg |
|  | agctccgttttcacggaaaa |
|  | ctacatgggaacagcctatt |
|  | caaaagaaggcacagaggcc |
|  | ttggtcaccaaatcagcatt |
|  | agaggctcagcaatgagtga |
|  | gggcgattacagacacaact |
|  | ctttatttctcgccactgaa |

**Table S2.**

**RT-PCR and qRT-PCR primers**

| **Reverse Transcription-PCR (RT-PCR)** | |
| --- | --- |
| **Primer name** | **Sequence** |
| F-IFNg_Ex1 | CTGTTACTGCCAGGACCCAT |
| F-IFNg_Int1 | GGGGCAGTATTTTATAGTGGGG |
| R-IFNg_Ex2 | GTTCCATTATCCGCTACATCTGA |
| F-IFNg_Int2 | AAGCTGAATATTCCCATTTGGC |
| R-IFNg_Ex3 | CAGCTTTTCGAAGTCATCTCGT |
| F-IFNg_Ex3 | ATGCAGAGCCAAATTGTCTC |
| R-IFNg_Ex4 | GCTTCCCTGTTTTAGCTGCT |
| F5-IFNg_Int3 | TGACCATCATGACATTAGCAGA |
| R3-IFNg_Ex4 | CTCTTCGACCTCGAAACAGC |
| F-TNFa_Ex1 | TGCTTGTTCCTCAGCCTCTT |
| F2-TNFa_Int1 | AAGGAGAGAGATGGGGGAGA |
| R-TNFa_Ex2 | GGCCAGAGGGCTGATTAGAG |
| F-TNFa_Int2 | GGTTTGGGGGTAGGGTTAGT |
| R-TNFa_Ex3 | TGGGCTACAGGCTTGTCACT |
| F-TNFa_Ex3 | GACAAGCCTGTAGCCCATGT |
| F-TNFa_Int3 | GCACAGGCCTTAGTGGGATA |
| R2-TNFa_Ex4 | AGGCCCCAGTTTGAATTCTT |
| **Real Time or Quantitative PCR (RT-qPCR)** | |
| IFNg RT2.0 FW | AGCTCTGCATCGTTTTGGGTT |
| IFNg RT2.0 RV | GTTCCATTATCCGCTACATCTGAA |
| TNFa Fw | GTTCCATTATCCGCTACATCTGAA |
| TNFa RV | TCAGCCTCTTCTCCTTCCTG |
| IL2 Fw | CAAGAATCCCAAACTCACCAG |
| IL2 RV | CGTTGATATTGCTGATTAAGTCC |
| 18S_3F | GTGGAGCGATTTGTCTGGTT |
| 18S_3R | AACGCCACTTGTCCCTCTAA |

**TableE S3. RL-PAT primers**

| **RNA ligation-mediated poly(A) test (RL-PAT)** | |
| --- | --- |
| PAT-anchor | 5’-rApp GGT CAC CTT GAT CTG AAG ddC-3’ |
| PAT-R1 | 5’-GCT TCA GAT CAA GGT GAC CTT TTT-3’ |
| PAT IFNy 3'UTR F | GGTTGTCCTGCCTGCAATATTTG |
| PAT TNF 3'UTR F | GGAGGACGAACATCCAACCTTC |

**Movie S1**

3D model of acquisition field of view generated from T-cell smFISH with 61 Z-stacks. The video shows multiple Teff cells on coverslips. Blue: nucleus (DAPI), green: *IFNG* mRNA (CALFluorRed 610), magenta: *TNF* mRNA (Quasar-670). In the left corner, scale bar: 5μm, adjusting according to movie magnification.

**Movie S2**

3D model of T-cell smFISH in three Teff cells. The video shows subsequentially staining for blue: nucleus (DAPI), green: *IFNG* mRNA (CALFluorRed 610), magenta: *TNF* mRNA (Quasar-670). Last image: merged. Scale bar: 5μm.
